# Imaging Nanoscale Nuclear Structures with Expansion Microscopy

**DOI:** 10.1101/2021.05.04.442164

**Authors:** Emma L. Faulkner, Jeremy A. Pike, Ruth M. Densham, Evelyn Garlick, Steven G. Thomas, Robert K. Neely, Joanna R. Morris

## Abstract

Commonly applied super-resolution light microscopies have provided insight into subcellular processes at the nanoscale. However, imaging depth, speed, throughput and cost remain significant challenges, reducing the numbers of three-dimensional, nanoscale processes that can be investigated and the number of laboratories able to undertake such analysis. Expansion microscopy solves many of these limitations but its application to imaging nuclear processes has been constrained by concerns of unequal nuclear expansion.

Here we demonstrate the conditions for isotropic expansion of the nucleus. Using DNA damage response proteins, BRCA1, 53BP1 and RAD51 as exemplars we quantitatively describe the three-dimensional nanoscale organisation of over 50,000 DNA damage response structures. We demonstrate the ability to assess chromatin regulated events and show the simultaneous assessment of four elements. This study thus provides the means by which expansion microscopy can contribute to the investigation of nanoscale nuclear processes.

## Introduction

Major processes central to life occur within eukaryotic nuclei such that high-resolution imaging of nuclear structures is critical to improving our understanding of DNA replication, DNA repair, gene regulation and transcription. The application of fluorescence microscopy has provided insight into the organisation and regulation of many of these processes and studies applying super-resolution microscopy (SRM) have allowed investigation of the spatial organisation of proteins within sub-compartments with nanoscale resolution.

A particular example is DNA damage signalling, where repair proteins are redistributed in a spatiotemporally regulated manner to form microscopically visible aggregates known as “foci” around damaged sites^1^. Application of confocal microscopy and SRM such as stimulated emission depletion (STED) microscopy, stochastic optical reconstruction microscopy (STORM), and structured illumination microscopy (SIM) have contributed to a spatial map of repair signalling in which a protein’s time of arrival and departure, and its relative site of residence directs DNA repair pathways^1–7^.

However, established super-resolution techniques offer a compromised solution to imaging of three-dimensional spatial organisation of nanoscale protein arrangements, typically requiring a trade-off between resolution and throughput^8^. Our current super-resolution view of DNA damage signalling, for example, is based on analysis of hundreds of structures with relatively low resolution, e.g. using SIM with a spatial resolution of 100-130 nm laterally^6^, or tens of structures with improved resolution e.g. STED, SMLM with lateral resolutions of 30-80 nm and 20 nm, respectively^3,5,7^. Moreover, because established SRM technologies require expensive equipment and sophisticated analytical tools, the number of laboratories able to investigate nanoscale structural organisations with the requisite sub-diffraction limit resolution is restricted.

Expansion Microscopy (ExM) has the potential to overcome some of the problems posed by other SRM modalities. However, whilst ExM has been successfully applied to the analysis of cytoplasmic structures, concerns over differential nuclear expansion and controversy on how samples should be prepared for its investigation has limited ExM investigation of nanoscale structures in the nucleus^9–14^.

Herein we demonstrate conditions for the isotropic expansion of the nucleus of human epithelial cells with minimal nanoscale distortion. We investigate the nanoscale organisation of DNA-damage response proteins 53BP1 and BRCA1, which have previously been assessed by established SRM techniques^3,4,6^, and use manipulation of chromatin regulators underpinning 53BP1 localisation to demonstrate the ability of ExM to assess chromatin-regulated events. We assess thousands of nanoscale nuclear features, enabling unprecedented description of substructure heterogeneity and illustrate 3D and four-colour analysis. These data demonstrate that ExM can be applied for the nanoscale analysis of nuclear structures at scale, offering an unparalleled insight into nuclear processes that is accessible to many laboratories.

## Results

### Isotropic Expansion of the Nucleus

A key consideration in applying ExM is to avoid anisotropic expansion that can result in sample distortion^15,16^. In ExM, specimens are labelled with conventional fluorescent antibodies or proteins equipped with anchors that enable their incorporation into dense and even polyelectrolyte gel meshwork formed throughout the sample. The sample is digested with proteases and the addition of water results in volumetric expansion of the gel, with the aim of retaining the relative spatial organisation of the labels^9,10,17^. However, the presence of genomic DNA in the gel has been suggested to introduce distortions when expanding the nucleus, adversely affecting isotropic expansion^12,14,18^.

We hypothesised that an approach in which nucleic acids are anchored into the gel might both maintain the relative spatial organisation of nuclear structures, many of which relate to nucleic acid processing, and also promote isotropic expansion of the nucleus. To test this, we employed a nucleotide alkylating agent, conjugated to an acryolyl group through NHS-ester chemistry, to form a compound termed ‘LabelX’^19^. This compound anchors polynucleotides into the gel network but its impact on the nanoscale structure of the nucleus in ExM is unknown.

We first examined nuclei in cells that had been grown in the presence of the thymidine analogue 5-ethynyl-2’deoxyuridine (EdU) to visualise the DNA via conjugation of a fluorescent azide^20^. Following the application of the ExM protocol, we noted nuclear areas were increased ~ 16x (an expansion factor of 4x in one dimension) and nuclear volumes increased by ~ 52x (an expansion factor of 3.7x in one dimension) (Supplementary Figure 1 a-d). We found that nuclear expansion measurements corresponded to the macroscale expansion of the gel (Supplementary Figure 2).

To assess the isotropy at the nanoscale the same nuclei were imaged pre- and post-expansion and features within these images were compared (Figure 1a). Axial expansion of the sample changes the imaging depth of field. Nevertheless, we were able to confirm that the morphology of the nuclei was retained, with the same nanoscale features identified pre-expansion readily observable post-expansion with no distortions evident (Figure 1b-c). These data suggest that, in the presence of the LabelX anchor, the nucleus expands isotropically on the micro- and nanoscale with minimal distortions to the genomic architecture.

**Figure 1.**
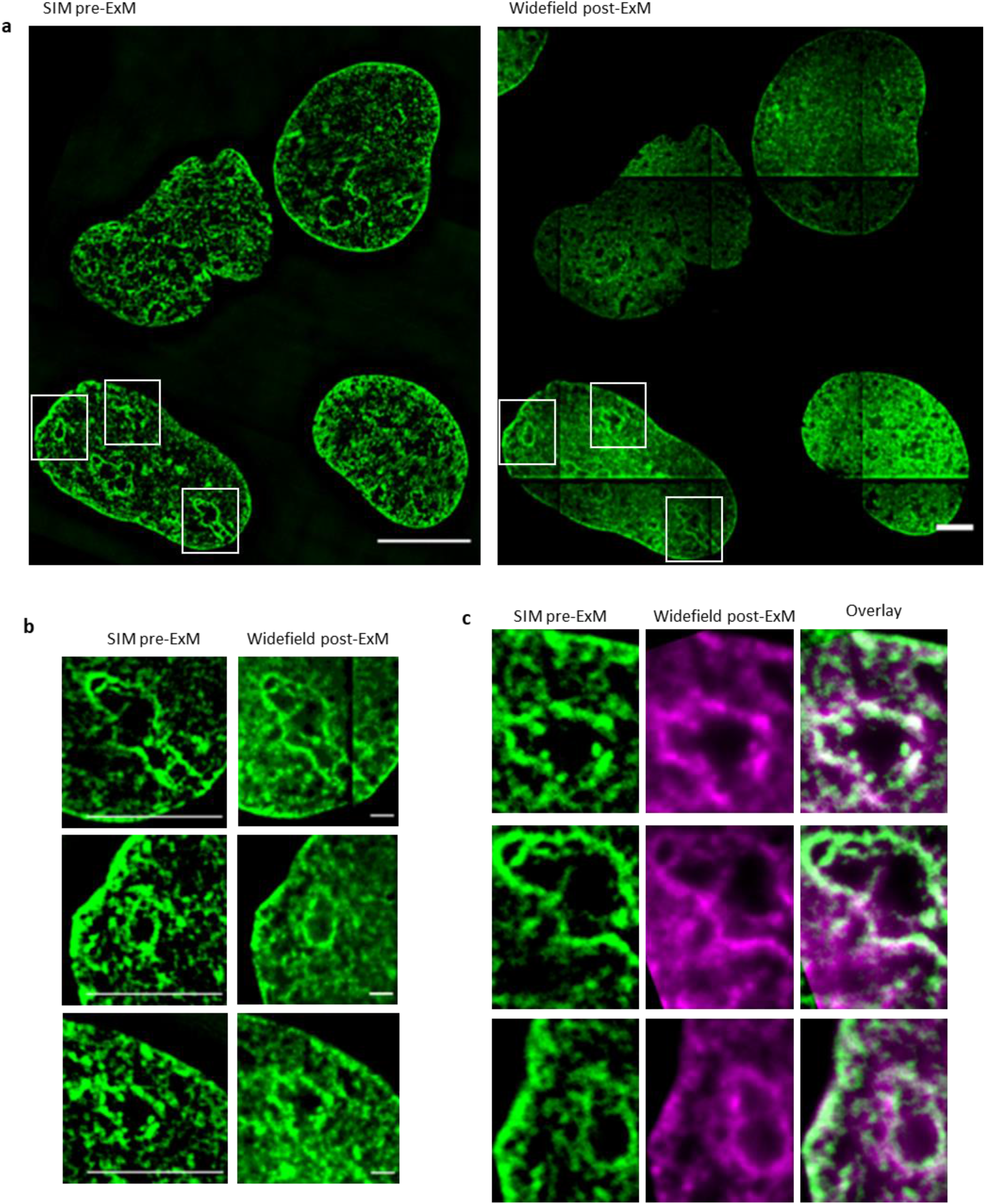
Isotropic expansion of the nucleus. U2OS cells were treated with EdU overnight prior to fixation. Click it reaction was performed for detection of EdU and images were acquired either pre-ExM or post-ExM. a) Correlative imaging was performed by acquiring pre-ExM images on a structured illumination microscope in 3D-SIM mode and then acquiring images of the same nuclei post-ExM on a widefield microscope. Scale bars 20 μm (equivalent to ~5 μm pre-ExM). b) Features were selected from the pre- and post-ExM correlative images and compared. Scale bars 5 μm. c) Corresponding features in the pre-ExM SIM (green) and post-ExM widefield (magenta) images were identified by eye and registered. Overlays of these features are shown.

### Nanoscale organisation of DNA-damage signalling proteins

We next investigated the organisation of pivotal DNA double-strand break repair regulator proteins, BRCA1 and 53BP1 in S-phase cells following exposure to gamma irradiation (2 Gy). The relative spatial organisation of these proteins under such conditions have previously been characterised by confocal and super-resolution techniques^2,4,6,7^.

Post-expansion 3D images were deconvolved and the organisation of 53BP1 and BRCA1 accumulations visually investigated (examples are shown in Figure 2a-c). Initial inspection suggested a heterogeneous population of 53BP1 and BRCA1 accumulations in early, mid and late S-phase classified nuclei. To quantitatively describe the spatial organisation of thousands of protein accumulations, we developed a semi-automated spot detection-based analysis which was applied to mid and late S-phase nuclei. The number of 53BP1 and BRCA1 spots within a 2 μm radius of a core BRCA1 spot was investigated. 9438 structures from 74 nuclei were characterised into 5 distinct classes (Supplementary Figure 3a-b) illustrating the ability of an ExM approach to capture and describe nuclear structure heterogeneity.

**Figure 2.**
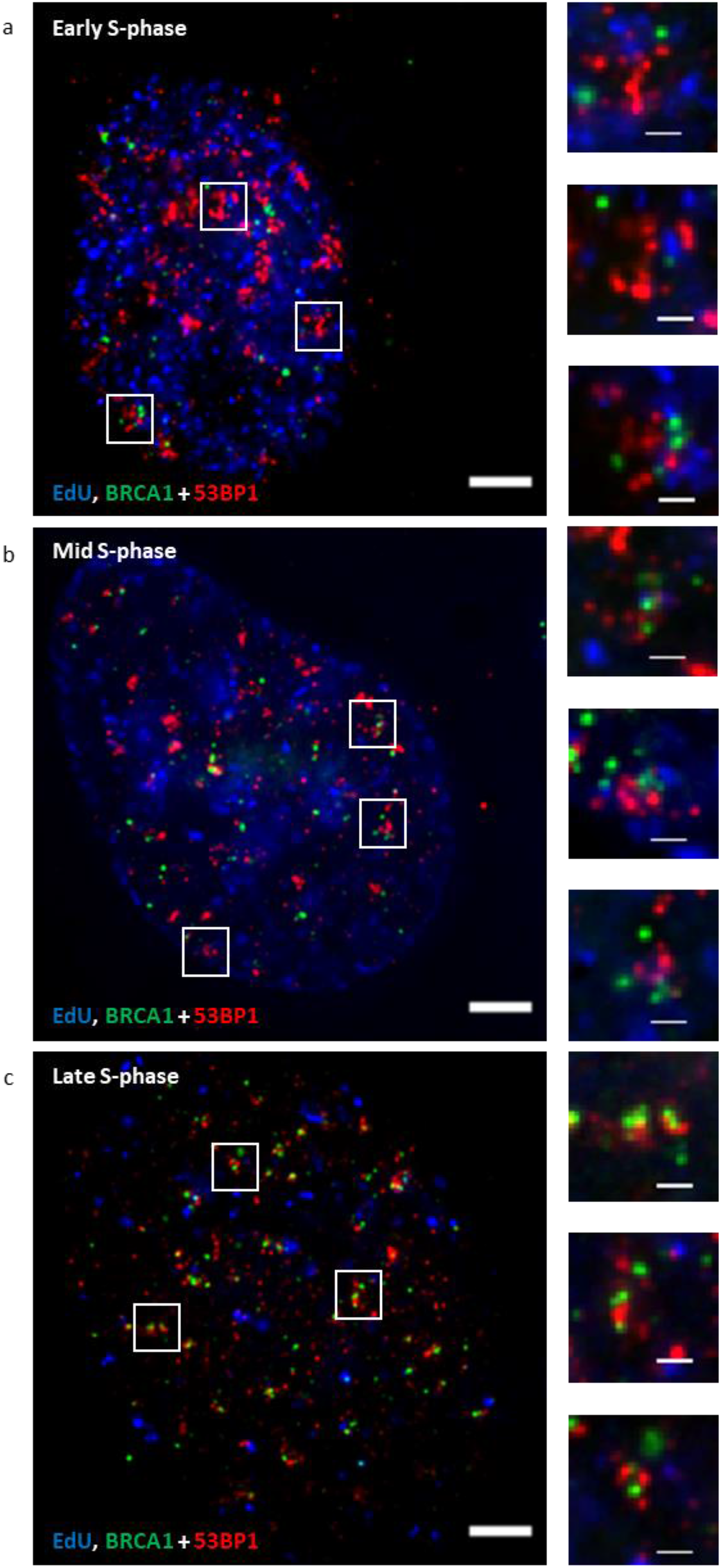
Imaging nanoscale organisation of DNA-damage signalling proteins, 53BP1 and BRCA1. U2OS cells were treated with EdU (blue) and damaged with irradiation (2 Gy) then allowed 1 hour to recover prior to fixation. Cells were immunostained for BRCA1 (green) and 53BP1 (red), then prepared using ExM method. Post-expansion images of nuclei classified as early (a), mid (b) and late S-phase (c). Scale bars 10 μm (large image) and 2 μm (selected regions), equivalent to ~2.5 μm and ~500 nm pre-ExM.

### Averaging of hundreds of nanoscale features to explore chromatin regulators

If ExM is to be routinely used for the examination of nanoscale nuclear structures, it must be capable of reporting on changes to protein distribution regulated by chromatin reorganisation. Previous observations have suggested that 53BP1 accumulations are influenced by BRCA1^2,6,21,22^ and by chromatin changes mediated by SMARCAD1 and the ubiquitin-specific protease 48 (USP48)^23,24^. SMARCAD1 has been reported to promote the localisation of 53BP1 to the periphery of irradiation (IR)-induced foci, whereas USP48 restricts 53BP1 positioning in BRCA1-proficient cells.

We treated cells with siRNA to SMARCAD1 or USP48 and subjected them to ExM, staining for BRCA1 and 53BP1 (Supplementary Figure 4a-b). By eye, 53BP1 appeared to occupy smaller volumes in SMARCAD1 depleted cells and larger volumes in USP48-depleted cells (Supplementary Figure 4c-d). 27,458 BRCA1:53BP1 structures from 279 nuclei from these experiments were assigned to one of the five defined classes. In SMARCAD1 depleted cells there were no significant changes in distributions of structures in mid-S-phase cells, whereas in late S-phase cells, fewer structures in which 53BP1 as a series of discontinuous spots surrounding a central BRCA1 (class 5) structures were observed compared to controls (Supplementary Figure 5a-b). In contrast, in USP48 depleted cells, we observed an increase in the number of class 5 structures in both mid and late S-phase cells (Supplementary Figure 5c-d). Intriguingly, class 5 structures resemble 53BP1:BRCA1 accumulation patterns previously described by confocal and SIM analysis in irradiated S-phase cells^2,6^ and to some 53BP1 structures in pre- and post-replicative cells^7^ (examples shown in Supplementary Figure 6). We investigated the average distribution of 53BP1 accumulations relative to the central BRCA1 spot in control cells by selecting over 100 examples of class 5 structures where proteins were oriented parallel to the focal plane and an average structure profile was generated. In the averaged class 5 profiles, the core BRCA1 spot spanned approximately 800 nm (equivalent to 200 nm pre-expansion). 53BP1 had a continuous localisation encapsulating BRCA1, spanning 2.5-3 μm diameter (equivalent to 625-750 nm pre-expansion). The peak-to-peak distance of the 53BP1 distribution was measured as 1.27 μm and 1.43 μm (equivalent to 0.32 μm and 0.36 μm pre-expansion) in mid and late S-phase structures, respectively (Figure 3a). In the orthogonal views of the averaged class 5 structure profiles, BRCA1 and 53BP1 occupy distinct regions with no visible overlap between them (Figure 3b). From averaged class 5 structure profiles, we estimate the separation between 53BP1 and BRCA1 to be equivalent to 60-80 nm, pre-expansion.

**Figure 3.**
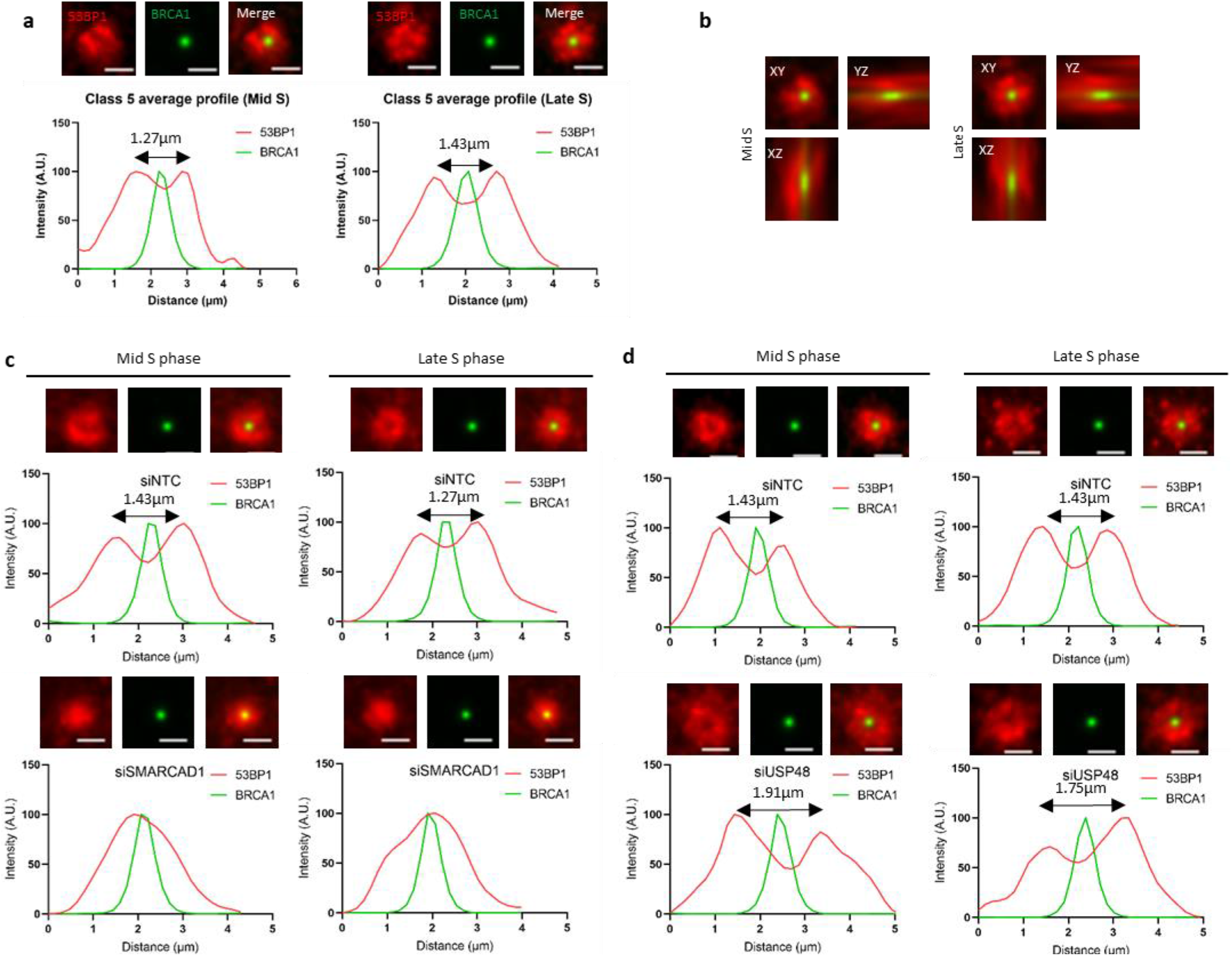
Positioning of BRCA1: 53BP1 following depletion of chromatin regulators SMARCAD1 and USP48. a) U2OS cells were treated as in Figure 2. Examples of class 5 structures were selected and an average profile generated. Scale bars 2 μm (equivalent to ~500 nm pre-ExM). N=3 (mid S = 130 structures, late S = 125 structures). b) Orthogonal views of average class 5 structures as in Figure 3a are shown. c) U2OS cells were treated with NTC or SMARCAD1 siRNA for 72 hours. Cells were treated with EdU (not shown) and damaged with irradiation (2 Gy) then allowed 1 hour to recover prior to fixation. Cells were immunostained for BRCA1 (green) and 53BP1 (red), then prepared using ExM method. Average class 5 structures shown in the image (red-53BP1, green –BRCA1, and then merged) and average intensity profiles generated from those structures are shown in the graph below, n=3 (NTC mid S = 143 structures, NTC late S = 184 structures, SMARCAD1-depleted mid S = 164 structures and SMARCAD1-depleted late S = 221 structures). Scale bars 2 μm (equivalent to ~500 nm pre-ExM). d) U2OS cells were treated as in c but USP48 siRNA was used. Average class 5 structures shown in the image (red-53BP1, green –BRCA1, and then merged) and average intensity profiles generated from those structures are shown in the graph below n=3 (NTC mid S = 72 structures, NTC late S = 77 structures, USP48-depleted mid S = 93 structures and USP48-depleted late S = 135 structures). Scale bars 2 μm (equivalent to ~500 nm pre-ExM).

To examine more closely the influence of chromatin regulators of 53BP1 using ExM, examples of tens of class 5 structures where both BRCA1 and 53BP1 were oriented approximately parallel to the focal plane were selected and averaged structure profiles were generated from cells treated with siRNA targeting SMARCAD1 or USP48 (Figure 3c-d, Supplementary Figure 7a-b). In the SMARCAD1 depleted cells, the void between the proteins was lost and 53BP1 occupied a smaller volume, consistent with the observation that 53BP1 has a peak intensity coinciding with that of BRCA1 in the absence of SMARCAD1^23^. In contrast, following USP48 siRNA treatment, a clear void was visible between the two proteins and the 53BP1 peak-to-peak distance measuring 1.92 μm and 1.75 μm (equivalent to 0.48 μm and 0.44 μm pre-expansion) compared to control values of 1.43 μm (equivalent to 0.36 μm pre-expansion) in both mid and late S-phase average structures. Additionally, 53BP1 accumulations in the average structures were positioned further away from the core BRCA1, spanning ~ 5 μm as compared to ~ 4 μm in controls (equivalent to 1.25 μm and 1 μm). These measurements are similar to those in published work using confocal microscopy where the peak-to-peak distance of 53BP1 was measured to be 0.3 μm in control cells and 0.5 μm following USP48 depletion^24^.

As these measurements were made on selected examples of class 5 structures (where BRCA1 and 53BP1 were oriented approximately parallel to the focal plane), next, all class 5 structures of all orientations were investigated in 3D (a total of 3,249 structures). Each class 5 structure was defined by the distance between the core BRCA1 spot and the surrounding 53BP1 accumulations (Supplementary Figure 7c-d). Following SMARCAD1 depletion, we observed >70% of class 5 structures from both mid and late S-phase nuclei had a reduced distance (defined as a separation of <0.5 μm) between 53BP1 spots and the core BRCA1 spots compared to controls where >85% of class 5 structures exhibited a separation distance of 1.8-2 μm between 53BP1 spots and the core BRCA1 spot. Correspondingly, in USP48 depleted cells, we observed >80% of class 5 structures had an increased distance (defined as a separation of ~ 2-2.5 μm) between 53BP1 spots and the core BRCA1 spot, whilst in the controls >70% of class 5 structures had a separation distance of ~ 1.8-2 μm between 53BP1 spots and the core BRCA1 spot.

### Four colour ExM of nanoscale nuclear structures

An advantage of ExM over several other SRM methods is the ability to perform multi-colour imaging for the interrogation of several nanoscale features simultaneously. To test this capability, post-expansion 3D images of nuclei immunostained for BRCA1, RAD51 and 53BP1 were acquired (Figure 4 a). We examined cells treated with control or USP48 siRNA as USP48 loss is associated with increased DNA resection lengths, increased RAD51 numbers and intensity^24^ and indeed post-expansion 3D images of nuclei showed that RAD51 accumulations were visually larger (Figure 4b & 4c) and late S-phase cells had more structures classified as those with multiple RAD51 spots associated with multiple 53BP1 spots (Class 4; Supplementary Figure 8a-d). We developed a classification approach to describe the spatial organisation of 3,908 structures in which RAD51 was arbitrarily defined as the centre of repair foci, and the relative spatial organisation of 53BP1 and/or BRCA1 relative to this centre was described. Using this method, we identified ten classes of structures (Supplementary Figure 9a-b). Of these classes, three (classes 7, 8 & 9) contained RAD51, 53BP1 and BRCA1 (Supplementary Figure 9c). In mid and late S-phase, structures defined as >1 RAD51 spot associated with multiple 53BP1 spots (class 4) were increased in number following USP48 depletion, as were late S-phase structures defined as multiple RAD51 spots with BRCA1 spot(s) associated and encapsulated by multiple 53BP1 spots (class 9) (Supplementary Figure 9a-b). We further investigated class 4 and 9 structures to assess the prevalence of continuous RAD51 accumulations within them and found increased numbers of class 4 and 9 structures with continuous accumulations following USP48 loss (Supplementary Figure 10a-c). We also noted the location of BRCA1 in class 9 structures varied; in the many selected examples BRCA1 localised to one end of the RAD51 accumulation, whilst in other cases, BRCA1 was located at the centre of the RAD51 structure (Figure 4d). Taken as a whole our observations demonstrate ExM allows for quantitative description of the spatial organisation of multiple proteins within nanoscale nuclear structures.

**Figure 4.**
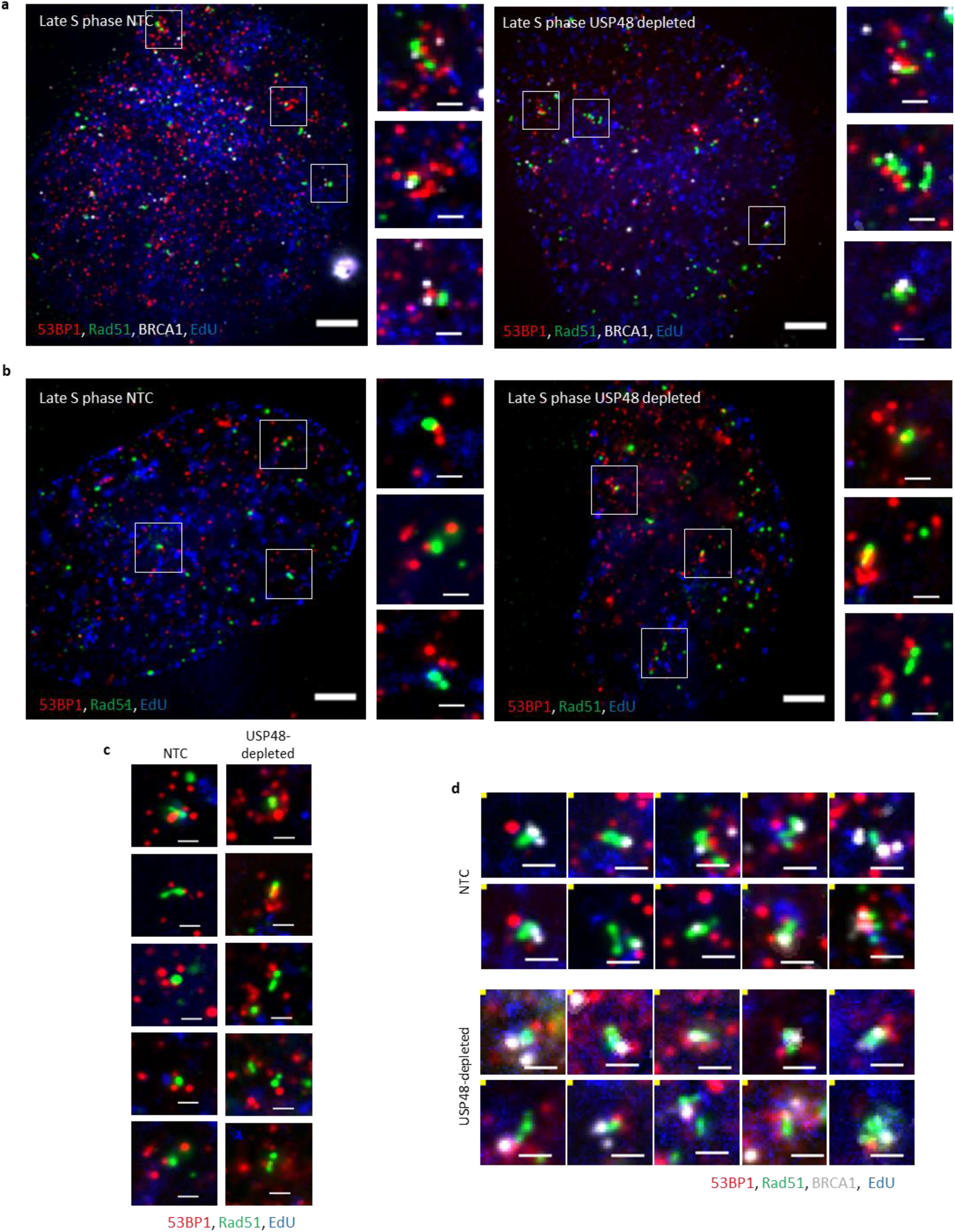
Visualising RAD51 accumulations. a) U2OS cells were treated with siNTC or siUSP48 for 72 hours. Cells were treated with EdU (blue) prior to irradiation (2 Gy) and allowed 1 hour to recover prior to fixation. Cells were immunostained for RAD51 (green), BRCA1 (white) and 53BP1 (red), then prepared using ExM method. Post-expansion images of late S-phase nuclei are shown. Scale bars 10 μm and 2 μm (equivalent to ~2.5 μm and 500 nm pre-ExM, respectively). b) U2OS cells were treated with siNTC or siUSP48 for 72 hours. Cells were treated with EdU (blue) prior to irradiation (2 Gy) and allowed 1 hour to recover prior to fixation. Cells were immunostained for RAD51 (green) and 53BP1 (red), then prepared using ExM method. Post-expansion images of late S-phase nuclei are shown. Scale bars 10 μm and 2 μm (equivalent to ~2.5 μm and 500 nm pre-ExM, respectively). c) Examples of co-enriched structures with RAD51 (green) and 53BP1 (red) from late S-phase nuclei are shown. Scale bars 2 μm (equivalent to ~500 nm pre-ExM). d) Examples of structures co-enriched with BRCA1 (white), RAD51 (green) and 53BP1 (red) from late S-phase nuclei are shown. Scale bars 2 μm (equivalent to ~500 nm pre-ExM).

## Discussion

Using the nucleic acid anchor, LabelX, and ~four-fold expansion we demonstrate retention of nuclear organisation in expanded polyacrylamide gel. The expansion factor of 3.7-4x in 1D was comparable to the macroscale expansion of the gel and features in pre-expansion SIM images and post-expansion widefield images were retained between the two acquisitions with no detectable distortions.

Additionally, our findings using ExM closely correlate with those of previous assessments of protein accumulations to damage sites using other super-resolution approaches. Firstly, the observation of a discontinuous spot-like appearance of 53BP1 similar to 53BP1-nanodomains visualised by STED microscopy^7^; secondly, our confirmation of the chromatin-mediated regulation of 53BP1 relationship with BRCA1, contracted by SMARCAD1 depletion and extended by USP48 loss, as previously described using confocal microscopy^23,24^; and finally, the observation of increased numbers of continuous RAD51 accumulations following depletion of USP48^24^. Thus, under our methodology, nanoscale changes driven by changes in chromatin regulation are readily detectable using ExM.

ExM prepared specimens are optically cleared, which eliminates the effect of light scattering throughout the sample and allows access to volumetric imaging on a conventional widefield microscope^25^. We used this aspect to assess the 3D spatial organisation of thousands of accumulations of DNA repair proteins in irradiated S-phase cells. The speed of volumetric image acquisition (widefield or SPIM) allowed large numbers of features to be captured and spot detection-based analysis used to describe spatial heterogeneity, here described in mid and late S-phase classified DNA-damage protein accumulation structures. This ability enables a complete overview of the distribution of structures without user bias.

The high throughput of ExM then allowed us to measure the average void between 53BP1 and BRCA1 in thousands of protein structures. Following SMARCAD1 depletion, the void between 53BP1 and BRCA1 was reduced/ absent, whilst following USP48 depletion the void between these proteins was increased. These structural changes were commensurate with those expected from previous observations^23,24^, indicating that ExM can be applied to faithfully detect nanoscale, chromatin-regulated changes within specific nuclear architecture.

While our aim in this study was to assess the suitability of ExM for measuring nanoscale features of the nucleus, some surprising observations herein lead to further questions. For example, whether the heterogeneous sub-populations of 53BP1 and BRCA1 co-enriched structures relate to repair of distinct types of DNA lesions/chromatin states^26^, and whether the single BRCA1 spot often observed at one end of discontinuous RAD51 structures represents loading of RAD51 from one side of the DNA break. Further application of ExM offers the ability to investigate these novel findings and to establish the inter-relations with chromatin and to other critical repair proteins.

ExM methodology has evolved rapidly, extending the numbers of biomolecules that can be labelled (e.g. lipids, sugars) and the types of labels used^9,10,27–29^. Its resolution has been improved by combining ExM with other SRM techniques^30–32^, by increasing expansion factor^33,34^ and by post-expansion labelling of biomolecules, which can improve the fidelity in the final image^35–37^. These have the potential to improve further the analysis of nanoscale nuclear features^38^. Thus, with the current methods in hand, ExM can contribute significantly to a quantitative understanding of nuclear processes.

## Acknowledgements

The authors would like to acknowledge the help of Dee Kavanagh (COMPARE), Iain Styles (University of Birmingham), Krystian Ubych (University of Birmingham), Alexandra Walker (University of Birmingham) and James Beesley (University of Birmingham) for advice in development of the project. JRM and RMD are supported by the University of Birmingham. The authors would like to acknowledge EPSRC (UKRI), grant numbers EP/L016346/1 ((EPSRC Centre for Doctoral Training in Physical Sciences for Health (Sci-Phy-4-Health)) and EP/N020901/1 for funding. SGT received funding from the British Heart Foundation (NH/18/3/33913) and COMPARE.

## Author Contributions

E.L.F performed ExM preparation of samples, image acquisition, image processing, associated data analysis and wrote the paper. J.A.P. wrote the groovy and Matlab scripts for the spot detection-based description of protein accumulations, and the generation of average structure profiles, respectively. R.M.D. helped in the validation of isotropic nuclear expansion and interpretation of data. E.G. performed pre-expansion SIM imaging and the registration and deformation determination of pre- and post-expansion images in the correlative imaging experiment. S.G.T co-supervised E.L.F and helped with data interpretation. R.K.N. and J.R.M co-supervised E.L.F., contributed to data interpretation, directed the project, and wrote the paper.

## Methods and Materials

### Antibodies and Reagents

A full list of siRNA sequences and antibodies can be found in Supplementary Tables 1–2. Western blots show representative images taken from more than three independent experiments unless otherwise stated. All chemicals unless otherwise stated are from Sigma or ThermoFisher.

### Cell lines

U2OS cells were grown in Dulbeccos Modified Eagle media (DMEM) supplemented with 10% fetal bovine serum (FBS) and 1% Penicillin/ streptomycin.

### Cell growth and EdU visualisation

U2OS cells were plated at a density of 5×10^4^ cells/ ml in a 24-well plate containing 13 mm #1.5 coverglass and treated with thymidine analogue, 5-ethynyl-2’deoxyuridine (EdU) (Life Technologies) at stated times at a concentration of 10 μM. Cells were then fixed with 4% PFA for 10 minutes at room temperature. Cells were permeabilised with 0.5% triton x-100 in PBS for 15 minutes. After blocking with 10% FBS in PBST for 20 minutes, EdU staining was carried out following Click-iT^®^ EdU Imaging Kits (Life Technologies) according to manufacturer’s instructions. Images were acquired prior to and following ExM preparation on widefield and selective plane illumination microscopes as stated.

### Radiation protocol

Immediately prior to irradiation, cells were treated with EdU at a final concentration of 10 μM. Cells were exposed to radiation with a Gamma-cell 1000 Elite irradiator (caesium-137 source) at a dose of 2 Gy. Cells were allowed to recover for 1 hour in DMEM supplemented with 10% FBS and 1% penicillin/ streptomycin.

### Correlative imaging

U2OS cells were treated in the same way as above. Following digestion of ExM treated samples, gels were cut into a distinctive shape for orientation of samples and images were acquired pre-expansion on a structured illumination microscope (SIM). Specimens were then expanded by addition of water and post-expansion images of the same nuclei were acquired on a widefield microscope.

### Transfections

siRNA transfections were carried out using the transfection reagent Dharmafect1 (Dharmacon), per the manufacturer’s instructions.

### Immunofluorescence staining and S-phase discrimination

Cells were subject to immunofluorescent staining prior to standard microscopy using a widefield microscope or were prepared using ExM methodology. Cells were plated at a density of cells 3×10^4^ cells/ ml in 24-well plate containing 13 mm #1.5 coverglass and treated as required. Cells were pre-extracted with CSK buffer (100 mM sodium chloride, 300 mM sucrose, 3 mM magnesium chloride and 10 mM PIPES, pH 6.8) for 1 minute at room temperature. Cells were fixed in 4% PFA for 10 minutes at room temperature and permeabilised with 0.5% Triton X-100 in PBS for 15 minutes at room temperature. After blocking with 10% FBS, EdU staining was carried out following Click-iT^®^ EdU Imaging Kits (Life Technologies). EdU incorporation can be detected by reaction with a fluorescent azide dye in a copper (I)-catalysed azide-alkyne cycloaddition^20^. EdU incorporation resulted in well-defined patterns of incorporation to discriminate between early, mid and late S-phase cells. Azide dyes used for EdU detection were AF488 (C10337, Life Technologies), AF405 (1307, Click Chemistry Tools) and AF647 (C10340, Life Technologies). Cells were incubated with primary antibodies at the stated concentrations for either 1 hour at room temperature or overnight at 4°C and with the secondary Alexa Fluor antibodies for 1 hour at room temperature (summarised in Supplementary Table 2).

### Sample Expansion

#### Anchor synthesis

Acryloyl-X (6-((acryloyl)amino)hexanoic acid, succinimidyl ester) (AcX) (A20770, ThermoFisher Scientific) was re-suspended in anhydrous DMSO with a final concentration of 10 mg/ml. This was then aliquoted and stored in a frozen desiccated environment for up to 2 months. Label-IT amine (MIR3900, Mirus Bio) (100 μg) was re-suspended in reconstitution solution (100 μl) with a final concentration of 1 mg/ml. 10μl of AcX was added to label-IT amine and reacted overnight with shaking at room temperature to produce LabelX. Subsequently, this was aliquoted and stored in a frozen desiccated environment for up to 2 months.

#### Anchoring of cellular DNA and proteins

Cells were washed with 20 mM MOPS pH7.7, and incubated with nucleic acid anchor, LabelX (at a final concentration of 0.006 mg/ml) in MOPS at 37°C overnight followed by two washes with 1xPBS. Following two washes with 1x PBS, cells were incubated with the protein anchor Acryloyl-X, (AcX) (0.1 mg/ml) in PBS for >6 hours at room temperature. Specimens were washed with 1xPBS before gelation.

### Gelation, digestion and expansion

Monomer solution (1xPBS, 2M NaCl, 8.625% (w/w) Sodium acrylate (97%, 744-81-3, Sigma Aldrich), 2.5% (w/w) acrylamide (79-06-1 Sigma Aldrich), 0.15% (w/w) N,N’-methylenebisacrylamide (110-26-9, Sigma Aldrich)) was mixed, frozen in aliquots and thawed prior to use. Concentrated stocks of ammonium persulfate (APS) (7727-54-0, Sigma Aldrich) and tetramethylethylenediamine (TEMED) (110-18-9, Sigma Aldrich) at 10% (w/w) in water were diluted into the monomer solution to concentrations of 0.2% (w/w) on ice prior to gelation, with the initiator (APS) added last. The gelation solution (80 μl) was placed on a parafilm-covered slide in a humidified chamber. Coverslips were inverted onto the droplet with cells face down. Gelation was allowed to proceed at 37°C for two hours in a humidified chamber. Gels were removed from the slide and immersed in digestion buffer (1xTAE, 0.5% Triton X-100, 0.8 M guanidine HCl) containing 8 units/ml and Proteinase K (P8107S, New England Biolabs Inc.) was added freshly to the digestion buffer. Gels were digested either at room temperature overnight or 37°C for 4 hours. The gels were removed from the digestion buffer and placed in 50 ml of water to expand. Water was exchanged every 30 minutes until expansion was complete (typically 3-4 exchanges).

### Expanded specimen handling for imaging

For 3D-SIM imaging, unexpanded gels were mounted on high tolerance #1.5 Ibidi glass bottomed dishes (Thistle Scientific, IB-81158). For widefield imaging, expanded gels were cut to fit in MatTek dishes with a glass coverslips of 35 mm diameter (MatTeK Life Sciences, P35G-1.5-14-C). Excess water was removed and gels were embedded in 2% low melting point (LMP) agarose to limit gel movement during image acquisition. For SPIM imaging, gels were cut to fit the SPIM holder and placed cell-side up in the SPIM holder. 2% LMP agarose was pipetted into the holder until the bottom of the holder was covered, taking care not to get agarose in the interface between the top of the ExM gel and the objective lens. Deionised water was then added to the SPIM holder containing the gel to fully immerse the gel for imaging (details below).

Post-expansion images of nuclei were acquired on a SPIM to enable good optical sectioning with minimal photo-damage to the specimen^39^. ExM prepared specimens are optically transparent due to the large amount of water absorbed by the polymer meaning the gels are refractively matched to the water immersion medium and objectives required by the SPIM^25^. These features minimised optical aberrations and minimal processing was required to visualise nanoscale features of the nuclear architecture.

### Image acquisition

Structured illumination microscopy (SIM) was performed on a Nikon N-SIM-S system (Ti-2 stand, Hamamatsu ORCA Flash 4.0 scientific CMOS dual cameras with Cairn splitter system, Nikon Perfect Focus, Chroma ET525/50m, ET595/50m, and ET 700/75m emission filters, Nikon laser bed with 405nm/488nm/561nm/640nm laser lines). A Nikon 100x 1.49 NA TIRF oil objective was used. NIS Elements v5 software was used to control the system and acquire pre-expansion 3D-SIM images. Expanded samples were imaged on ASI RAMM microscope frame. Widefield imaging was performed using a Nikon 100× TIRF (N.A. 1.45) objective and an Evolve Delta EM-CCD camera, via a quad-band emission filter (Semrock, 432/515/595/730 nm). iSPIM was performed using twin Nikon 40x (N.A. 0.8) water-dipping objectives, a similar quad-band emission filter (Semrock, 432/515/595/730 nm) and a Hamamatsu ORCA Flash 4.0 scientific CMOS camera. Illumination for both setups was from a Cairn Research laser bank containing 100mW 405 nm, 150 mW 488 nm, 50 mW 561 nm and 100 mW OBIS 640 nm CW lasers. Light was directed to the sample via a quad-band dichroic mirror (Semrock, 405/488/561/635 nm). Micromanager was used to control the system and scan the sample.

### Image processing

Images acquired on the Nikon N-SIM-S system were reconstructed using stack reconstruction in the NIS elements software. Where stated, post-expansion image data was deconvolved using Huygens professional version 19.04 (Scientific Volume imaging, the Netherlands, http://svi.nl). A theoretical point spread function (PSF) was generated based on the microscope parameters and images were deconvolved using a classical maximum likelihood estimation (CMLE), a non-linear iterative restoration method which optimises the likelihood the objects in the estimated image are correctly localised based on the image and the PSF. This restoration method relies on the generation of an estimate of an object (synthetic image) which is compared to the measured image. The result of this is used to improve the original until the “difference” between the synthetic and measured image reach a minimum. Parameters for deconvolution were tested on example data sets for each experiment to determine optimal values, and these deconvolution templates were used for subsequent image processing and experimental repeats.

### Segmentation of nuclei

Quantification of nuclear areas pre- and post-expansion was performed using a script written in Matlab. Briefly, pre- and post-expansion images were processed by performing a rolling ball background subtraction and .tif files were saved into corresponding directories. Images were segmented using a manually determined threshold based on histograms generated for the images. Nuclei were segmented resulting in a mask of pixels where fluorescence is above the manually determined threshold. Boundaries were traced onto the binary image and then these boundaries were super-imposed onto the original image to determine the efficacy of segmentation. Centroids for each nucleus were determined and labelled. The areas of the labelled objects were calculated in pixels and microns^3^. Optimal parameters were determined and then used to determine nuclear areas pre- and post-expansion. A maximum and minimum area was defined based on the assumption that nuclei roughly conform to a circle, to remove features too small to be nuclei and features which correspond to >1 nucleus.

For determination of nuclear volumes pre- and post-expansion, images were segmented using auto-threshold in ImageJ to generate a mask. Otsu threshold efficiently segmented nuclei from background pixels. The generated mask was eroded and dilated, and any gaps were filled in. The volumes of the final mask was measured for each image. For each image, an object map was generated and compared to the original images to determine the efficacy of the segmentation.

### Registration of pre-ExM SIM and post-ExM images

For alignment and overlay of SIM and deconvolved expansion images in Figure 1c, corresponding features were identified by eye and registered with the FIJI plugin TurboReg (based on the work of Thévenaz et al., 1998) using a scaled rotation of the form x = λ { {cos θ, −sin θ}, {sin θ, cos θ} } ⋅ u + Δu. Landmarks were manually defined for the source (ExM) and target (SIM) images.

### Spot-detection based analysis of nuclear structures

Spot detection-based analysis of structures was performed using customised groovy script written for the open-source application Fiji^40^ available at https://github.com/JeremyPike/expansion-analysis. First spots were detected in each channel of interest by finding local maxima in a Laplacian of Gaussian filtered volume. Local maxima were filtered based on the prominence of each maxima relative to its local neighbourhood (spot quality). Spot detection was implemented using the TrackMate^41^ plugin. Spots in the assigned central site channel (e.g. BRCA1) were then clustered by finding the connected components of the graph formed by linking all spots within a fixed radius. Cropped 3D volumes centred on the centre of mass for each cluster were presented to the user to manually classify the type of repair foci structures. In this work, we analysed foci structures containing two or three repair proteins simultaneously.

General workflow for spot detection-based analysis of nanoscale nuclear structures:

1. All parameters were defined (summarised in Supplementary Table 3–4).
2. Spot detection was performed as described above according to parameters set and detections were displayed in 3D in the whole nucleus.
3. We determined whether an image was suitable for subsequent classification of structures. Images were omitted if sample movement or photo-bleaching was evident in the images.
4. Each feature (defined by the presence of a spot in the site channel) was displayed in the crop box which had a size of 5 μm^3^ and classified.

All experiments and subsequent spot detection-based analysis parameters are shown in Supplementary Table 4). In all experiments, cells were treated with EdU prior to irradiation (2 Gy) and allowed to recover for 1 hour.

### Generation of average foci structures and colocalisation analysis

To generate average structures of class 5 structures, examples of structures were selected where BRCA1 and 53BP1 were orientated parallel to the focal plane. These example structures were selected by visualising each class 5 structure in 3D using the spot detection based algorithm. The example structures were organised into directories, and then an average structure was generated using a customised Matlab script available at https://github.com/JeremyPike/expansion-analysis. The script generates the average structures as three-dimensional image stacks which were used to generate the maximum projections and orthogonal views. Additionally, the script generated a radial profile for each channel in the average structure. The radial profile was produced by binning pixels in the central slice into bands of varying distance (fixed width) from the site centre. The average intensity of all pixels in the band determines the radial profile at a specific distance.

### Defining the position of 53BP1 accumulations relative to core BRCA1 spot

To describe the placement of 53BP1 accumulations relative to the core BRCA1 spot following USP48 or SMARCAD1 loss, all class 5 structures for mid and late S-phase classified nuclei were sub-classified according to the distance of 53BP1 accumulations from the central BRCA1. These definitions are supplied in Supplementary table 5.

## siRNA sequences

**Supplementary Table 1.**
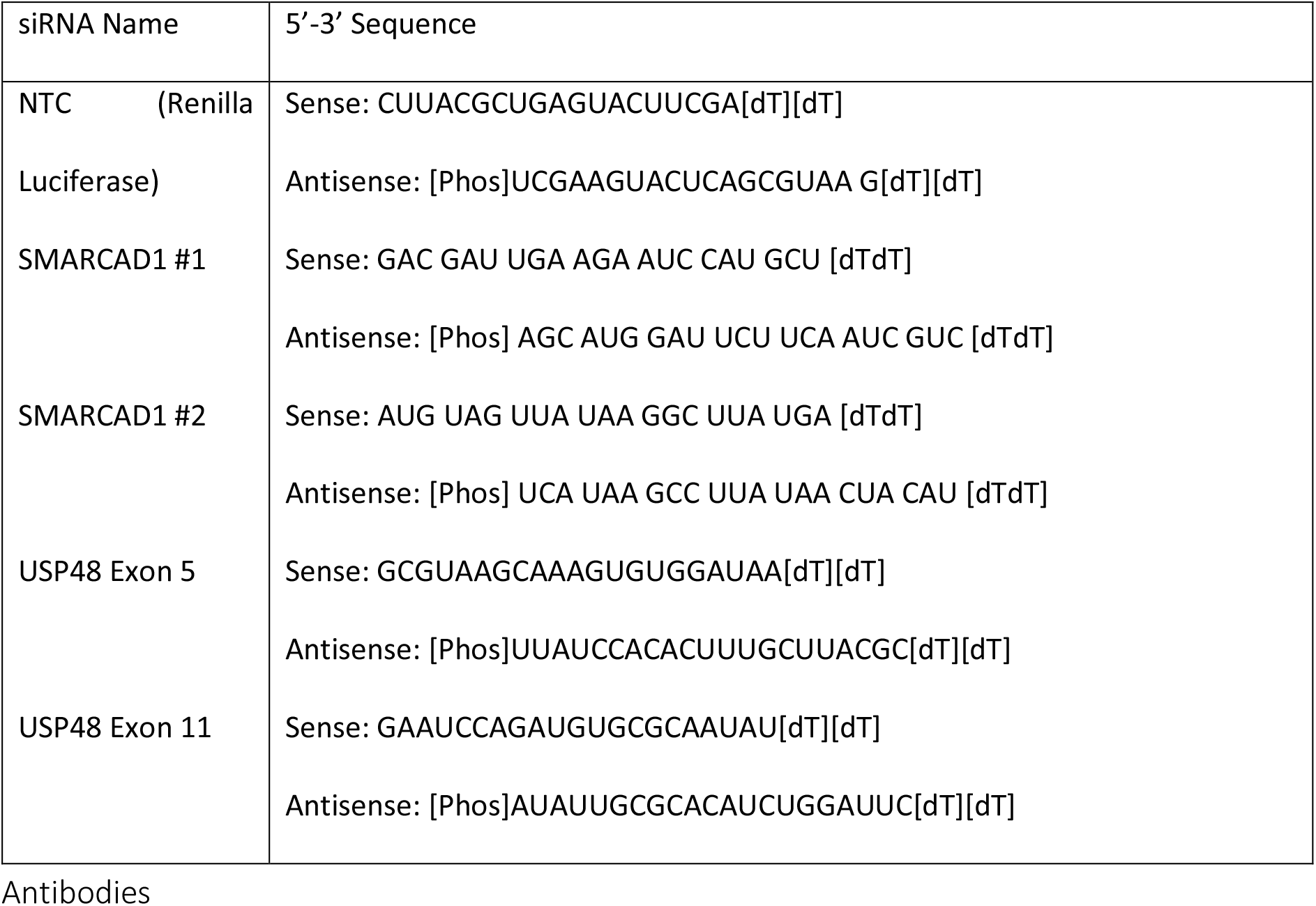
siRNA sequences.

**Supplementary Table 2.**
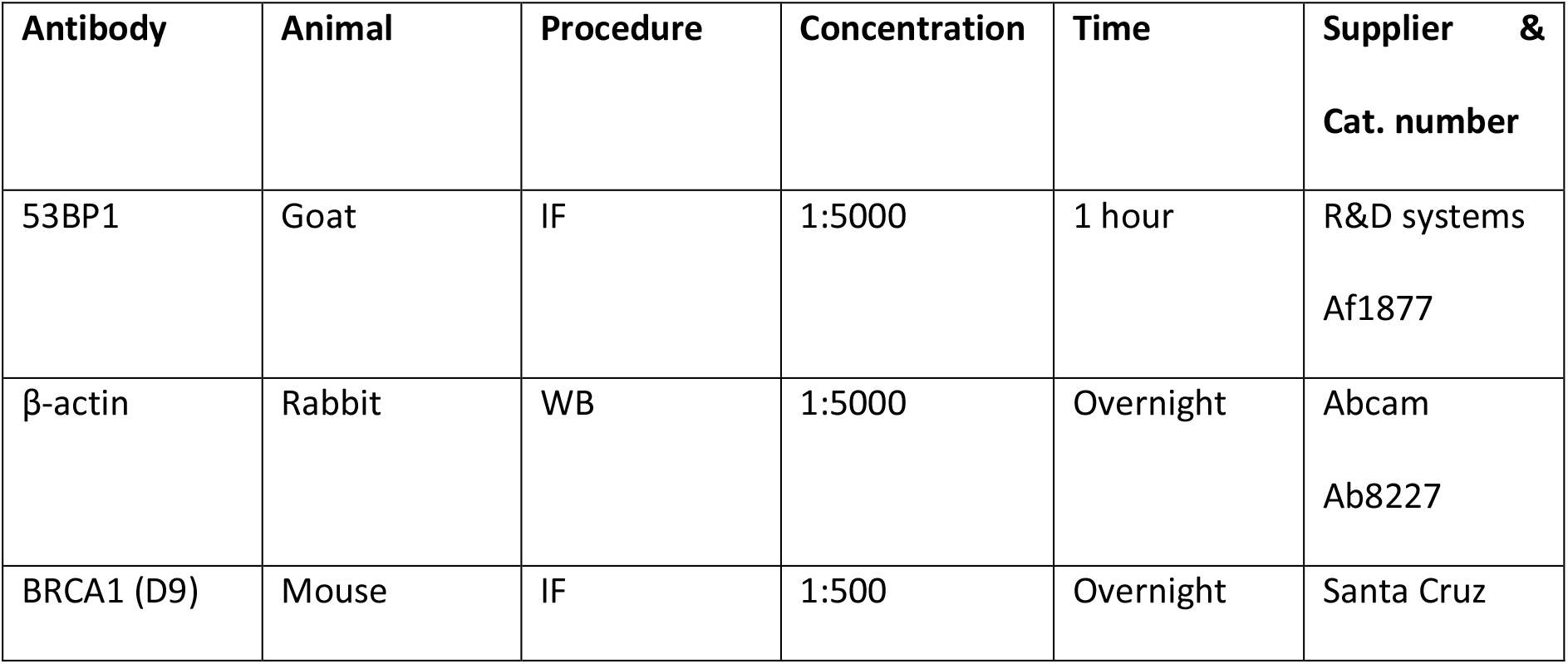
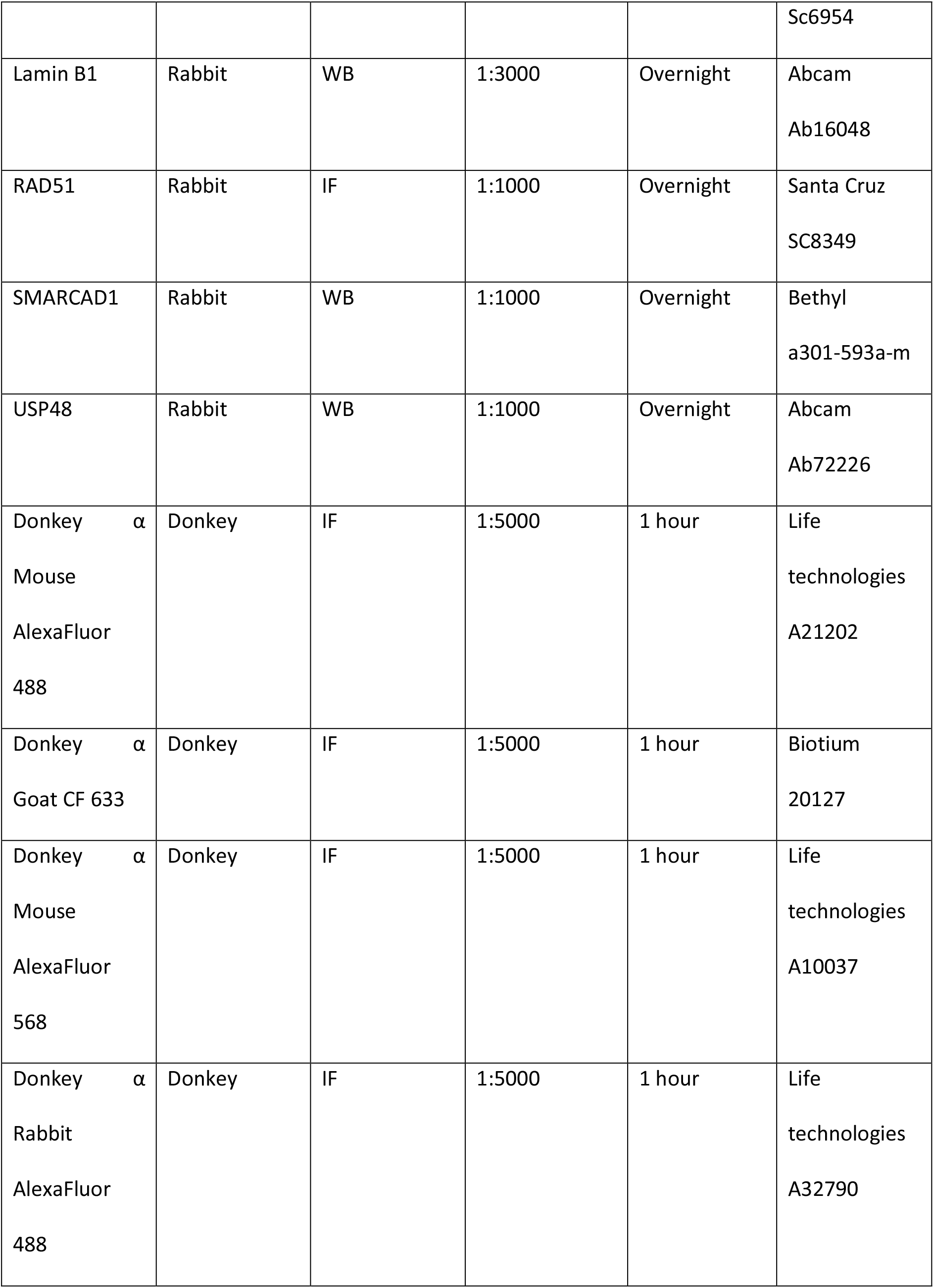
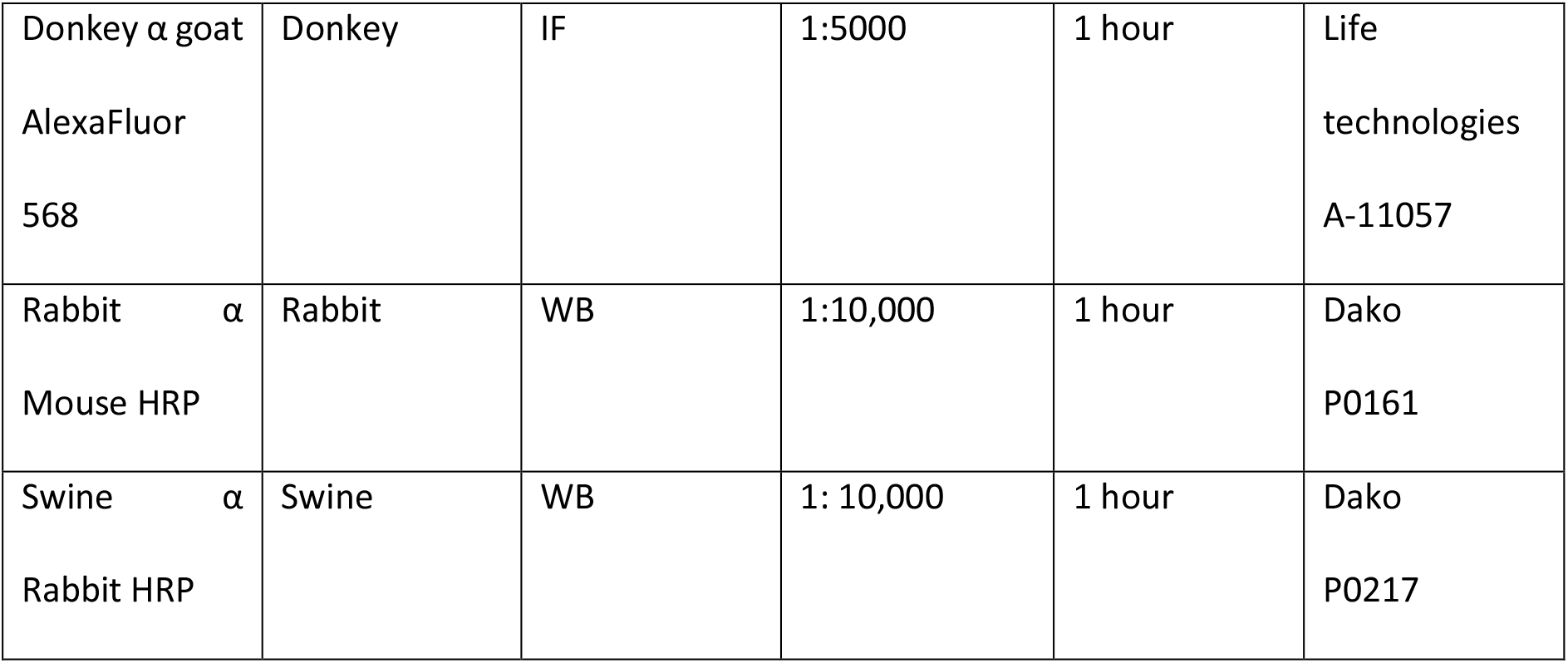
Antibodies including species raised in, concentration, conditions and protocols.

**Supplementary Table 3.**
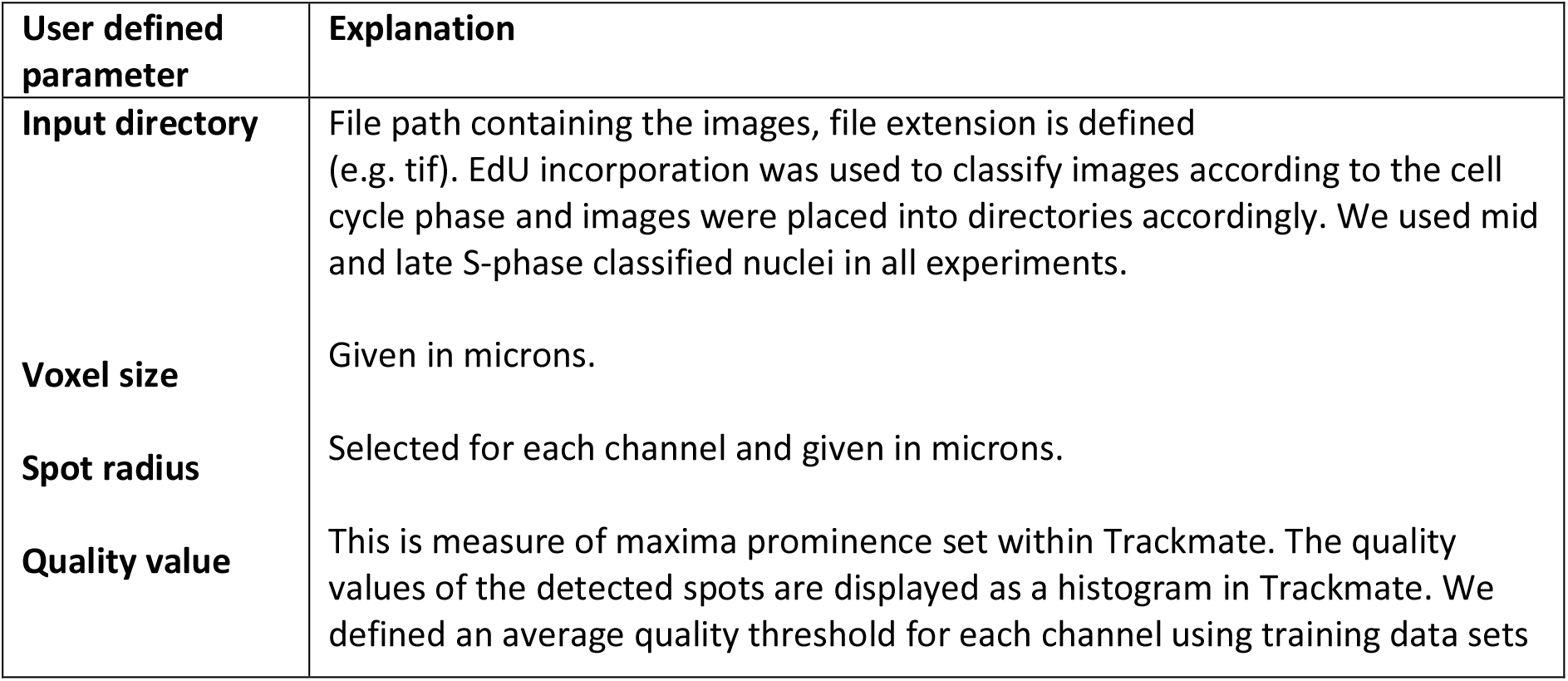
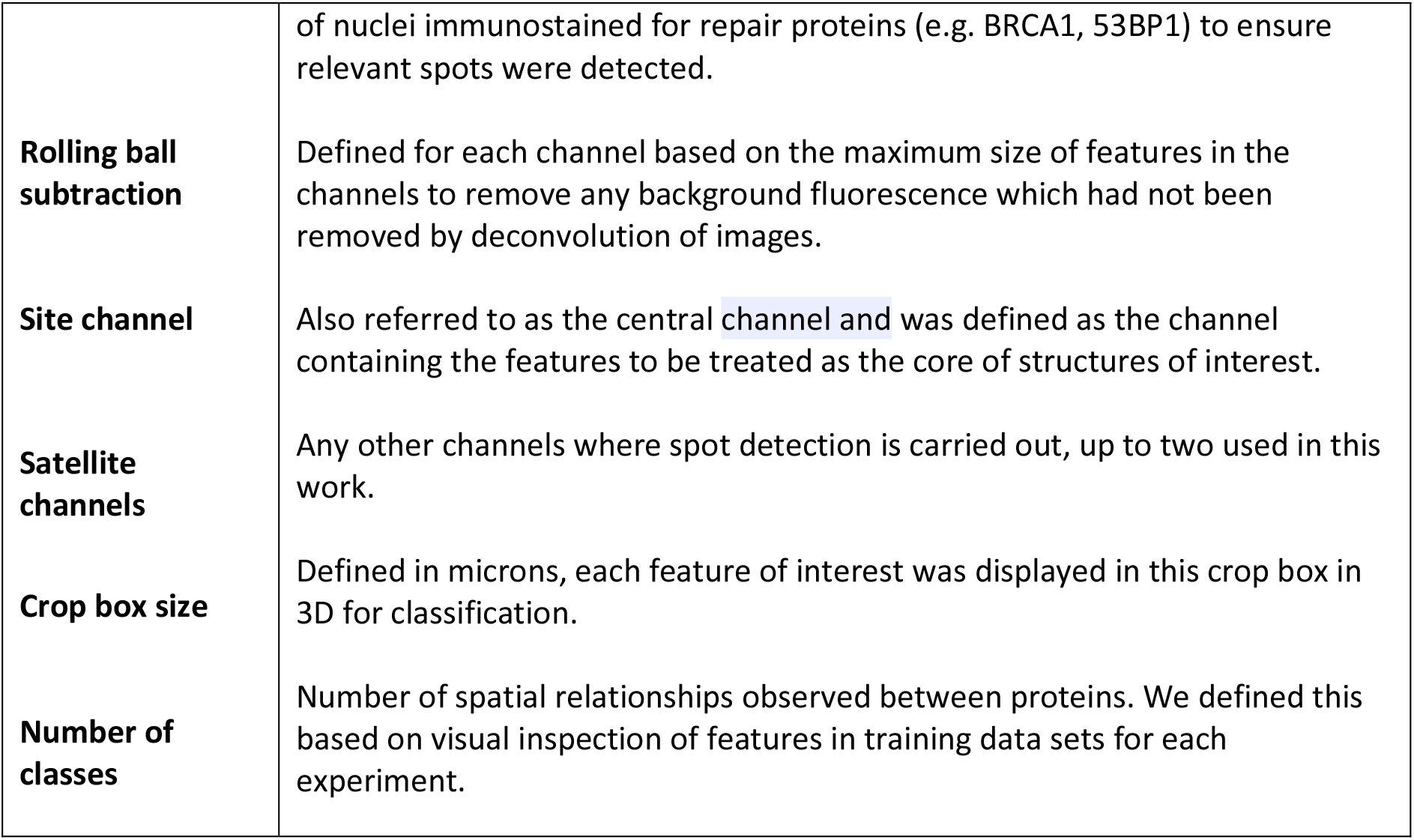
User defined parameters for spot detection-based analysis.

**Supplementary Table 4.** Summary of analysis parameters used to analyse BRCA1, 53BP1 and RAD51 accumulations in mid and late S-phase classified cells.

| Experiment* | siRNA | Primary antibodies | Spot detection-based analysis script | Central spot | Satellite spot(s) | Spot radius (μm) | Quality value | Rolling ball subtraction values (pixels) | Number of classes identified |
| --- | --- | --- | --- | --- | --- | --- | --- | --- | --- |
| Investigating BRCA1 and 53BP1 accumulations | N/A | BRCA1, 53BP1 | Two-colour analysis | BRCA1 | 53BP1 | 0.75 | 7.5 (53BP1)<br>6 (BRCA1) | 40 (53BP1)<br>14 (BRCA1) | 5 |
| Investigating BRCA1 and 53BP1 accumulations following SMARCAD1 and USP48 loss | siSMARCAD1<br>siUSP48 | BRCA1, 53BP1 | Two-colour analysis | BRCA1 | 53BP1 | 0.75 | 7.5 (53BP1)<br>6 (BRCA1) | 40 (53BP1)<br>14 (BRCA1) | 5 |
| Investigating RAD51 and 53BP1 accumulations following USP48 loss | siUSP48 | RAD51, 53BP1 | Two-colour analysis | RAD51 | 53BP1 | 0.75 | 7.5 (53BP1)<br>6 (RAD51) | 40 (53BP1)<br>14 (RAD51) | 6 |
| Investigating organisation of RAD51, 53BP1 and BRCA1 following USP48 loss | siUSP48 | BRCA1, 53BP1, RAD51 | Four-colour analysis | RAD51 | 53BP1<br>BRCA1 | 0.75 | 7.5 (53BP1)<br>6 (BRCA1, RAD51) | 40 (53BP1)<br>14 (BRCA1, RAD51) | 10 |

\*In all experiments, cells were treated with EdU (for cell cycle phase classification) prior to irradiation (2 Gy) and allowed 1 hour to recover.

**Supplementary Table 5.** Defining the positioning of BRCA1: 53BP1 following depletion of chromatin regulators, SMARCAD1 and USP48.

| **Definition of 53BP1 profile** | **Separation distance of 53BP1 from core BRCA1 ($\mu\text{m}$)** |
| --- | --- |
| Control | ~1.8-2 |
| Reduced | <0.5 |
| Increased | ~2-2.5 |

