## Supplemental Figures for "Imaging Nanoscale Nuclear Structures with Expansion Microscopy"

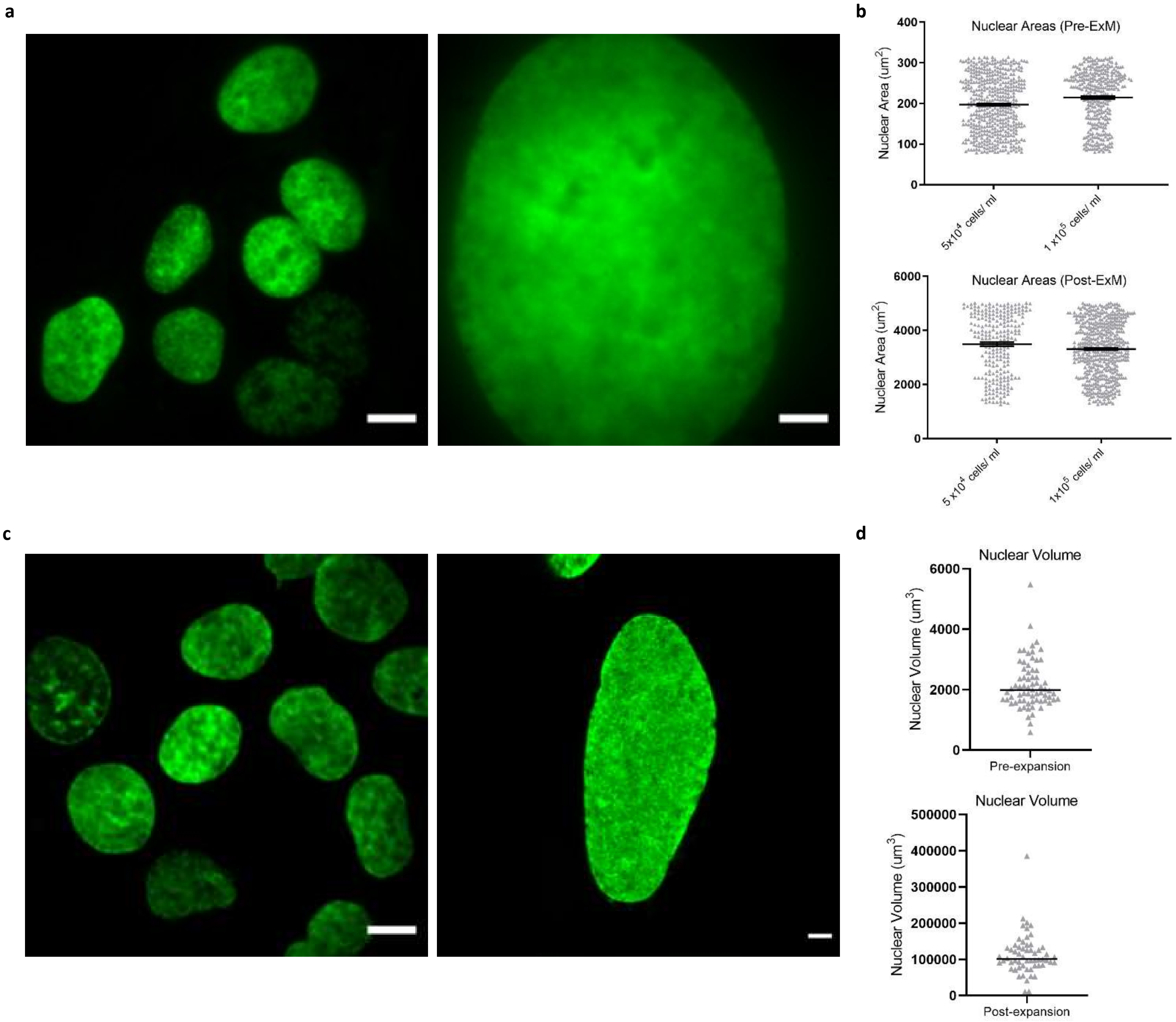


**Supplementary Figure 1. Isotropic Expansion of the nucleus.** U2OS cells were treated with EdU overnight prior to fixation. Click it reaction was performed for detection of EdU and images were acquired either pre-expansion or post-expansion.

a) 2D images of nuclei were acquired pre- and post-ExM. EdU is shown in green. Scale bars 10 µm.

b) Nuclear areas were quantified by segmenting nuclei from background pixels. Means ± s.e.m.

c) 3D images of nuclei were acquired pre-ExM on a widefield microscope and deconvolved. Post-ExM images of nuclei were acquired on a SPIM. Images are shown as maximum intensity projections. EdU is shown in green. Scale bars 10 µm and 40 µm, respectively.

d) Nuclear volumes were quantified pre- and post-ExM. Mean ± s.e.m.

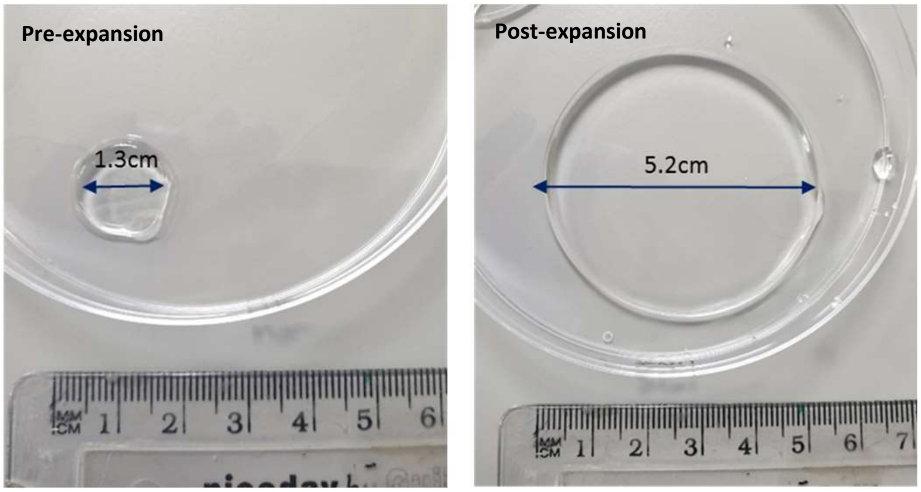


**Supplementary Figure 2.** The diameter of ExM gels was measured pre- and post-expansion.

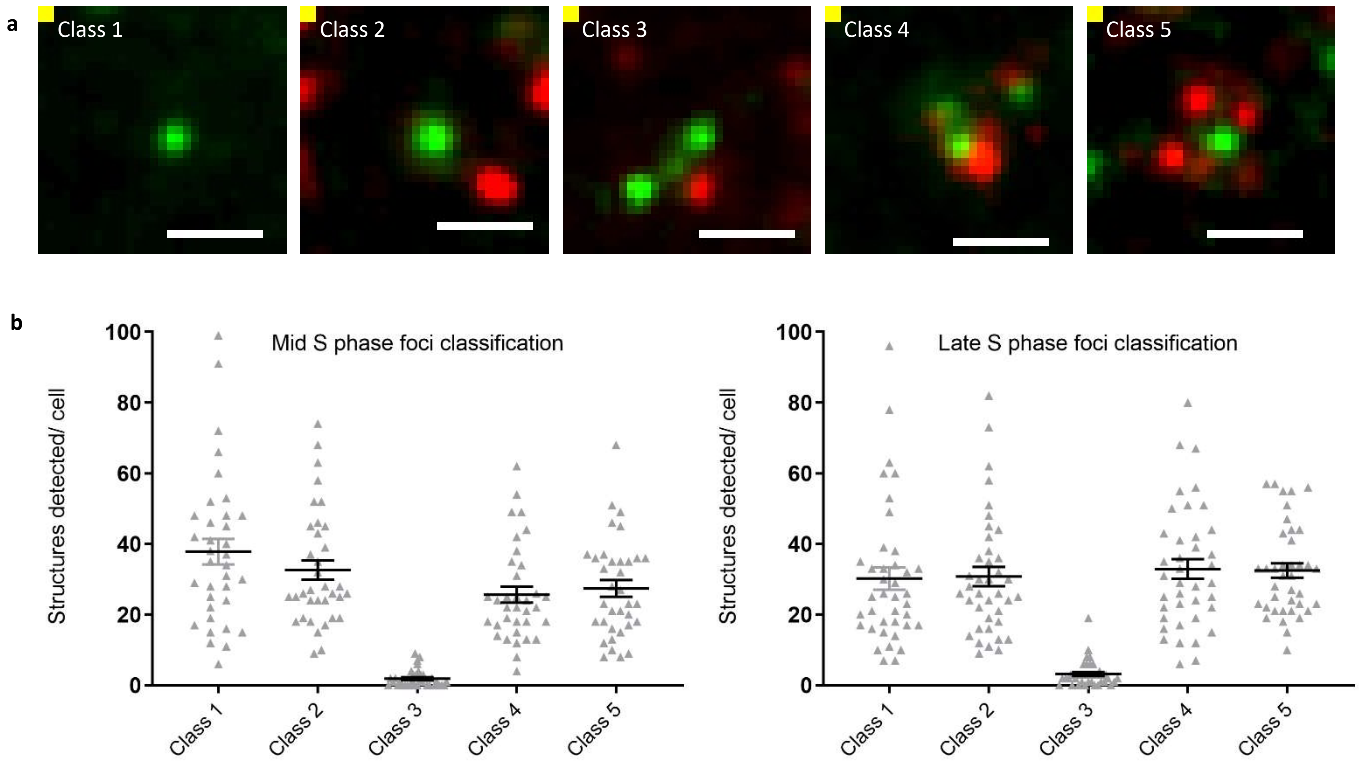


**Supplementary Figure 3. Characterising the nanoscale organisation of 53BP1 and BRCA1 co-enriched structures.**

a) Representative images of structure classes 1-5 containing BRCA1 spots (green) and 53BP1 spots (red). Scale bars 2 µm (equivalent to 500 nm pre-ExM). Class 1 structures were comprised of only the core BRCA1 spot. Class 2 structures contained the core BRCA1 spot and a 53BP1 spot. Class 3 structures contained multiple BRCA1 spots and one 53BP1 spot. Class 4 structures incorporated multiple 53BP1 and BRCA1 spots. Class 5 structures were defined as one BRCA1 spot encapsulated by multiple 53BP1 spots

b) Quantification of structure classes for mid and late S-phase classified nuclei. Means ± s.e.m. n=3, (mid S phase = 35 nuclei, 4387 structures & late S phase = 39 nuclei, 5051 structures).

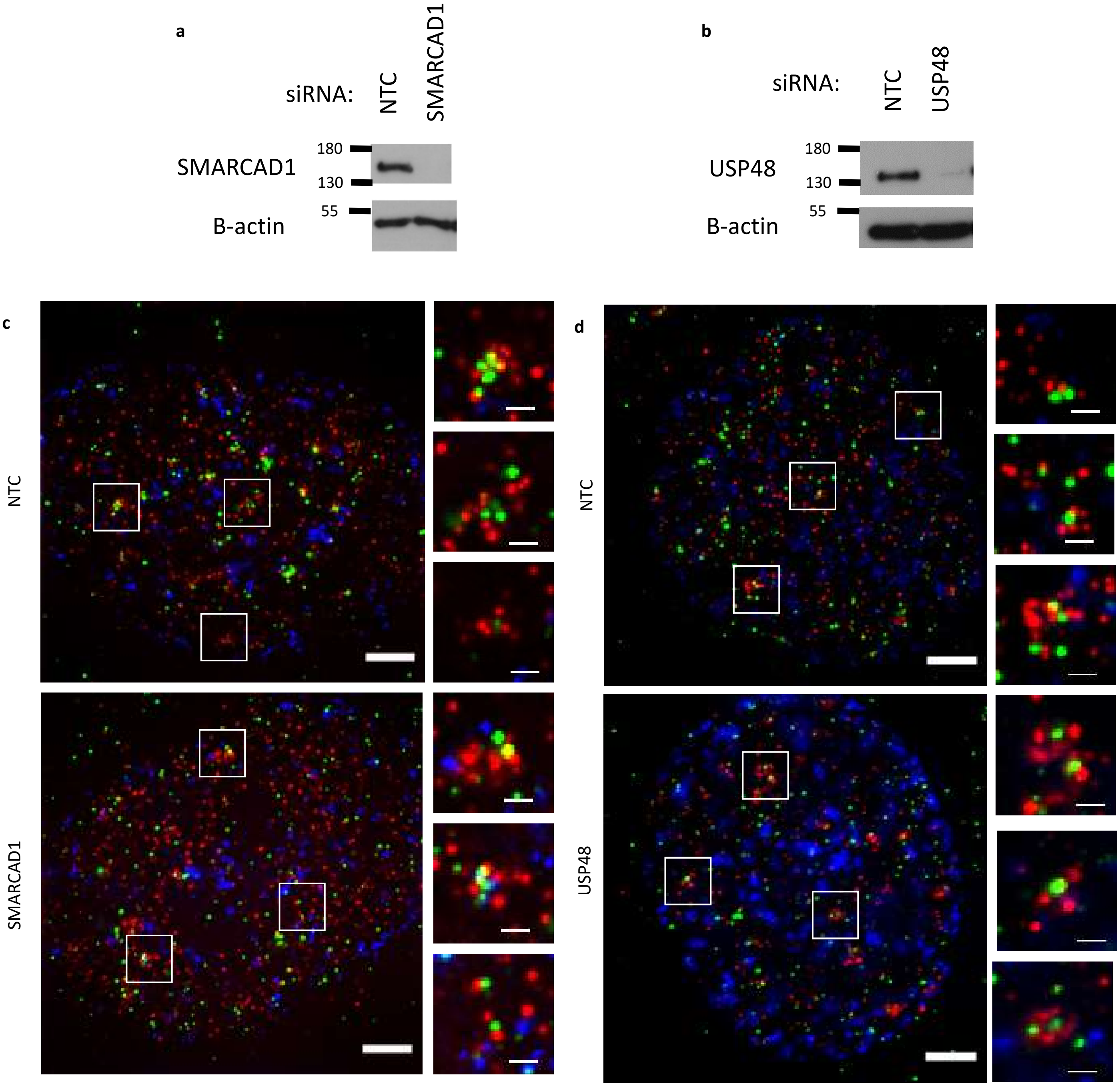


**Supplementary Figure 4. 53BP1:BRCA1 imaging following depletion of chromatin regulators, SMARCAD1 and USP48.**

a) U2OS cells were treated with siNTC or siSMARCAD1 for 72 hours prior to lysis. SMARCAD1 depletion was confirmed using antibodies shown.

b) U2OS cells were treated with siNTC or siUSP48 for 72 hours prior to lysis. USP48 depletion was confirmed using antibodies shown.

c) U2OS cells were treated with siNTC or siSMARCAD1 for 72 hours. Cells were treated with EdU (blue) for 1 hour prior to irradiation (2 Gy) and allowed 1 hour to recover prior to fixation. Cells were immunostained for BRCA1 (green) and 53BP1 (red), then prepared using ExM method. Post-expansion images late S-phase classified nuclei were acquired. Scale bars 10 µm (large images) and 2 µm (selected features), equivalent to ~2.5 µm and 500 nm pre-ExM, respectively.

d) U2OS cells were treated with siNTC or siUSP48 for 72 hours. Cells were treated with EdU (blue) for 1 hour prior to irradiation (2 Gy) and allowed 1 hour to recover prior to fixation. Cells were immunostained for BRCA1 (green) and 53BP1 (red), then prepared using ExM method. Post-expansion images late S-phase classified nuclei were acquired. Scale bars 10 µm (large images) and 2 µm (selected features) equivalent to ~2.5 µm and 500 nm pre-ExM, respectively.

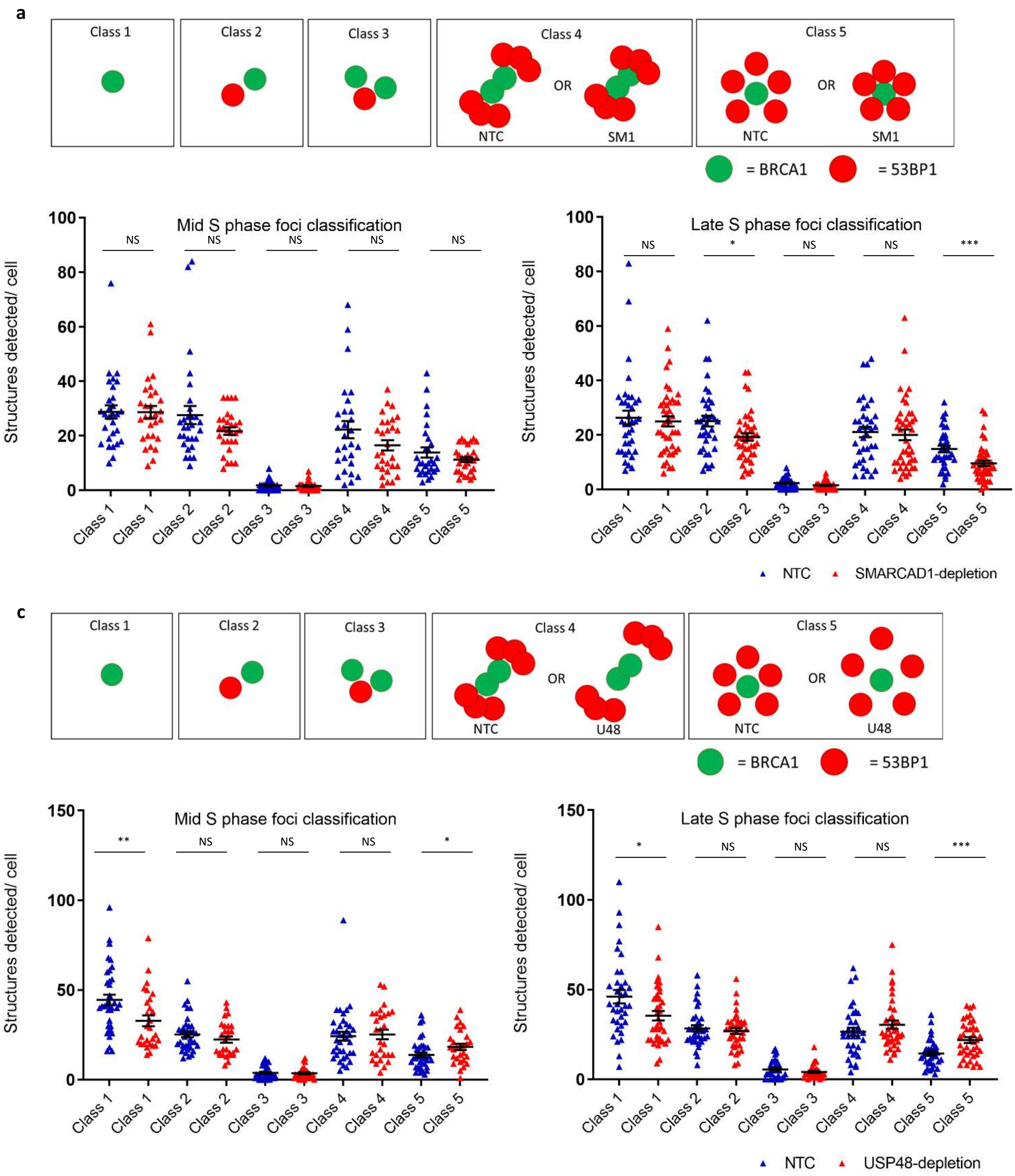


**Supplementary Figure 5**. **The spatial organisation of thousands of nanoscale BRCA1:53BP1 features after depletion of chromatin regulators SMARCAD1 and USP48.**

a) Schematic representation of structure classes 1-5.
b) Quantification of structure classes in mid and late S-phase cells following SMARCAD1 depletion (red) compared to controls (blue), n=3, >20 nuclei per sample. Means ± s.e.m. (NTC mid S = 29 nuclei, 2712 structures, NTC late S = 37 nuclei, 3320 structures, SMARCAD1 mid S = 29 nuclei, 2312 structures, SMARCAD1 late S = 43 nuclei, 3242 structures). Throughout Figure, ***P < 0.005; *P < 0.05 NS, not significant by two-tailed Student's t test.
c) Schematic representation of structure classes 1-5.
d) Quantification of structure classes in mid and late S-phase cells following USP48 depletion (red) compared to controls (blue), n=3, >20 nuclei per sample. Means ± s.e.m. (NTC mid S = 37 nuclei, 4133 structures, NTC late S = 36 nuclei, 4357 structures, USP48 mid S = 28 nuclei, 2867 structures, USP48 late S = 38 nuclei, 4515 structures). Throughout Figure, ***P < 0.005; **P <0.01;*P < 0.05 NS, not significant by two-tailed Student's t test.

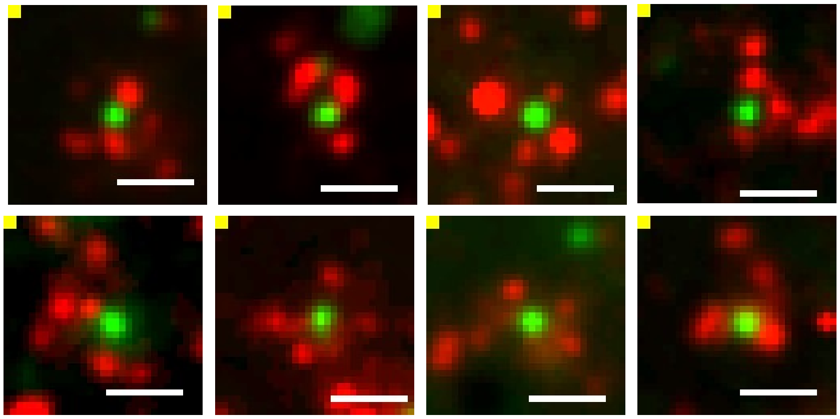

**Supplementary Figure 6.**Class 5 structures are comprised of spot-like accumulations of 53BP1 encapsulating the core BRCA1 spot. Examples of class 5 structures were selected from late S-phase classified nuclei as in Figure 2. Scale bars 2 µm, equivalent to 500 nm pre-ExM.

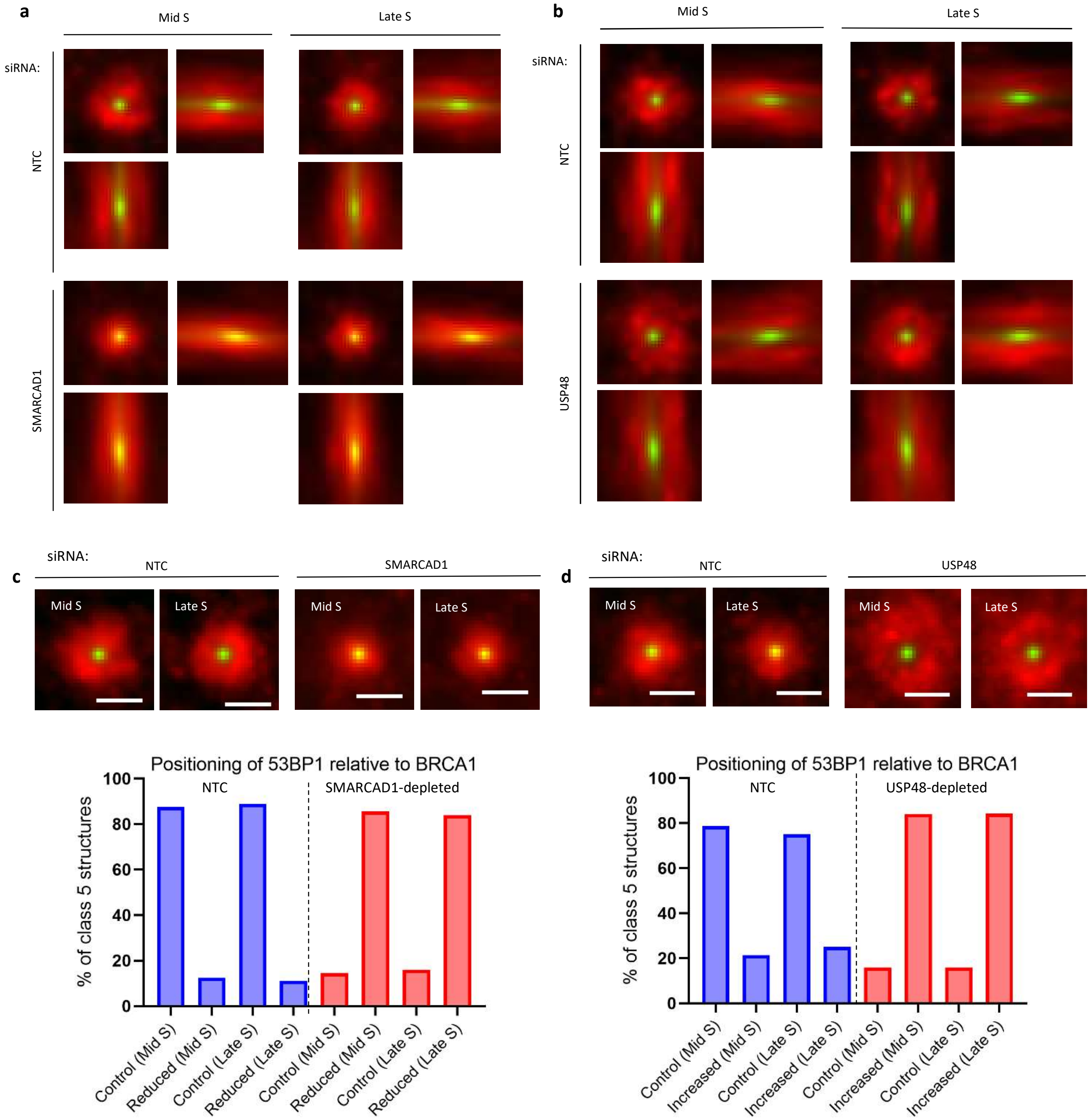


**Supplementary Figure 7. Positioning of BRCA1: 53BP1 following depletion of chromatin regulators SMARCAD1 and USP48.**

a) Orthogonal view of average class 5 structures for siSMARCAD1 and siNTC treated cells as in Figure 3c are shown. n=3. (NTC mid S = 143 structures, NTC late S = 184 structures, SMARCAD1-depleted mid S = 164 structures and SMARCAD1-depleted late S = 221 structures).

b) Orthogonal view of average class 5 structures for siUSP48 and siNTC treated cells as in Figure 3d are shown, n=3 (NTC mid S = 72 structures, NTC late S = 77 structures, USP48-depleted mid S = 93 structures and USP48-depleted late S = 135 structures).

c) All class 5 structures in siSMARCAD1 and siNTC treated cells were classified as having a reduced separation between BRCA1 and 53BP1 (defined as <0.5 µm) or a separation distance of ~1.8-2 µm (referred to as a control separation distance as determined in Figure 3a), respectively. Average class 5 structures were generated (shown as maximum projections, scale bar 2 µm), and the percentage of class 5 structures classified as having normal or reduced separation are shown. N=3 (NTC mid S = 360 structures, NTC late S = 469 structures, SMARCAD1-depletion mid S = 298 structures, SMARCAD1-depletion late S = 311 structures).

d) All class 5 structures in siUSP48 and siNTC treated cells were classified as having an increased separation between BRCA1 and 53BP1 of ~2-2.5 µm or a separation distance of ~1.8-2 µm (referred to as a control separation distance as determined in Figure 3a), respectively. Average class 5 structures were generated (shown as maximum projections, scale bar 2 µm) and the percentage of class 5 structures classified as having normal or increased separation are shown. N=3 (NTC mid S = 414 structures, NTC late S = 384 structures, USP48-depletion mid S = 404 structures, USP48-depletion late S = 609 structures).

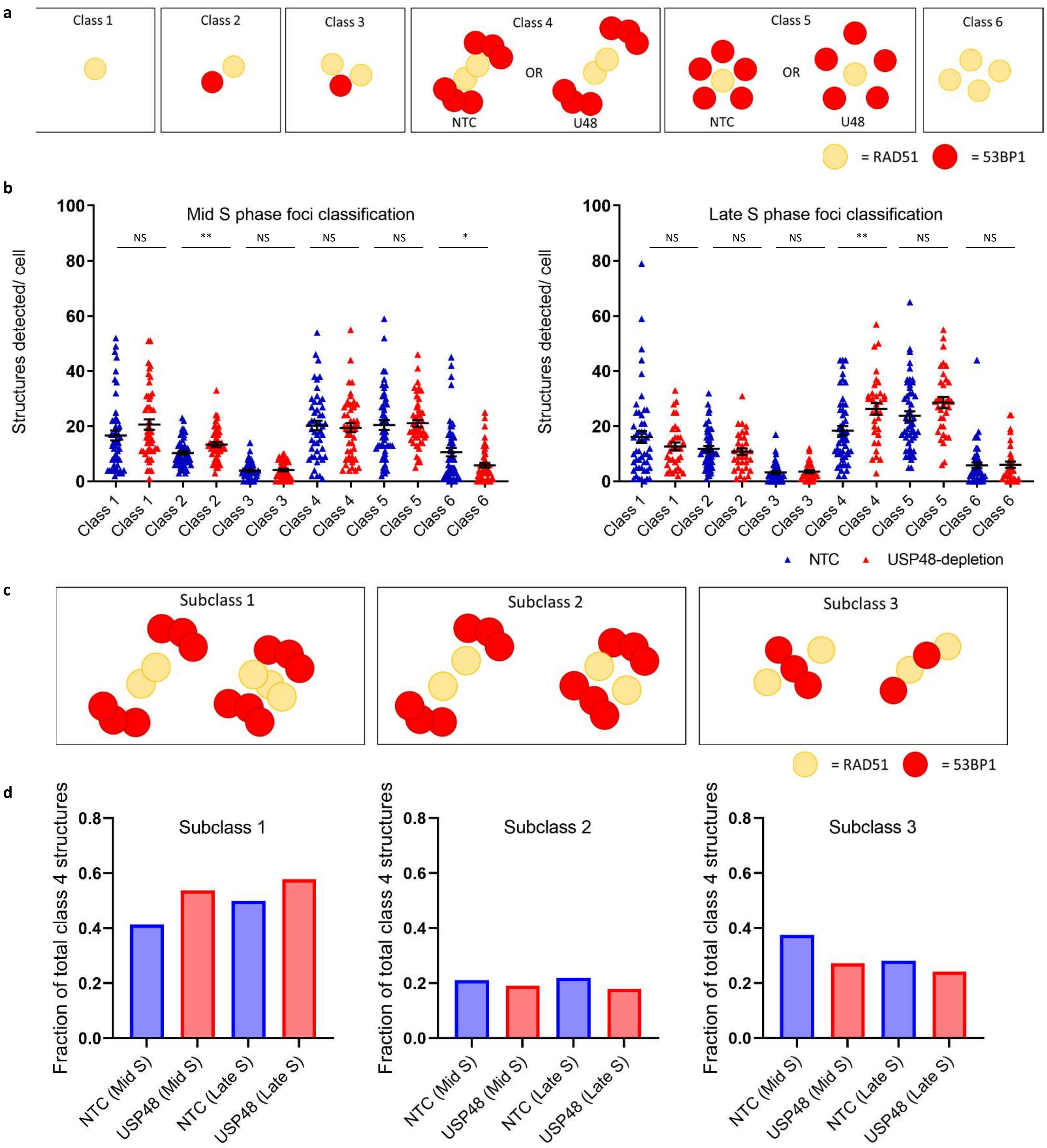


**Supplementary Figure 8. The spatial organisation of thousands of nanoscale RAD51:53BP1 features following USP48 depletion**.

a) Schematic representation of structure classes 1-6 with RAD51 (yellow) and 53BP1 (red).

b) Quantification of structure classes in mid and late S-phase cells in siUSP48 and siNTC treated cells, n=3, >20 nuclei per sample (NTC mid S = 51 nuclei, 4156 structures, USP48-depleted mid S = 54 nuclei, 4118 structures, NTC late S = 48 nuclei, 4054 structures, USP48-depleted late S = 34 nuclei, 2992 structures). Means ± s.e.m. Throughout Figure, **P <0.01;*P < 0.05 NS, not significant by two-tailed Student's t test.

c) Schematic representation of sub-classes in class 4 structures: subclass 1 (defined as continuous RAD51 structures encapsulated by multiple 53BP1 spots within a 2 µm radius), subclass 2 (defined as discontinuous RAD51 structures encapsulated by multiple 53BP1 spots within a 2 µm radius) or subclass 3 (defined as multiple RAD51 spots associated with multiple 53BP1 spots within a 2 µm radius).

d) All class 4 structures were sub-classified subclass 1, subclass 2 or subclass 3 as shown schematically. N=3, NTC mid S = 642 structures, NTC late S = 401 structures, USP48-depleted mid S = 518 structures, USP48-depleted late S = 756 structures.

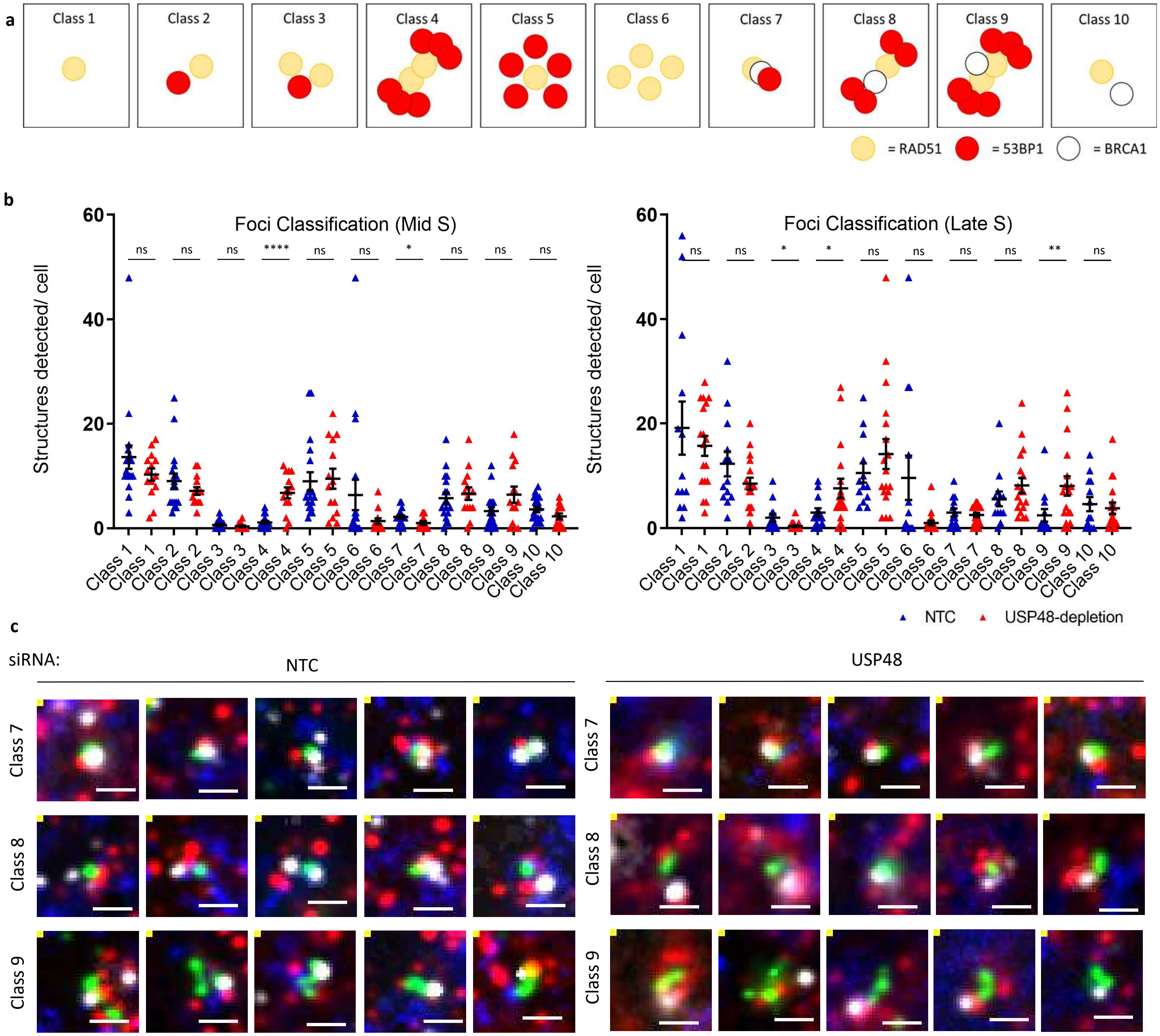


**Supplementary Figure 9. The spatial organisation of nanoscale RAD51, BRCA1 and 53BP1 features following USP48 depletion.**
U2OS cells were treated with either control or USP48 siRNA for 72 hours. Cells were treated with EdU (blue) prior to damage induction with irradiation (2 Gy) and allowed 1 hour to recover prior to fixation. Cells were immunostained for RAD51 (green), BRCA1 (white) and 53BP1 (red) and then prepared using ExM.

a) Schematic representation of structure classes 1-10 with RAD51 (yellow), BRCA1 (white) and 53BP1 (red).

b) Quantification of structure classes in mid and late S-phase cells following USP48 depletion compared to controls, n=2, >10 nuclei per sample. Means ± s.e.m. (NTC mid S = 19 nuclei, 983 structures, USP48-depleted mid S = 14 nuclei, 725 structures, NTC late S = 13 nuclei, 940 structures, USP48-depleted 1260 structures) Throughout Figure, **** P<0.001; **P <0.01;*P < 0.05 NS, not significant by two-sided Student's t test.

c) Examples of class 7 (defined as BRCA1, RAD51 and 53BP1 spots which are closely associated), class 8 (defined as 1 RAD51 spot and 1 BRCA1 spot encapsulated by multiple 53BP1 spots) and class 9 (defined as multiple RAD51 spots with BRCA1 spot(s) associated and encapsulated by multiple 53BP1 spots) structures selected from late S-phase classified nuclei are shown for USP48 depleted cells and controls. Scale bars 2μm (equivalent to ~500nm pre-ExM).

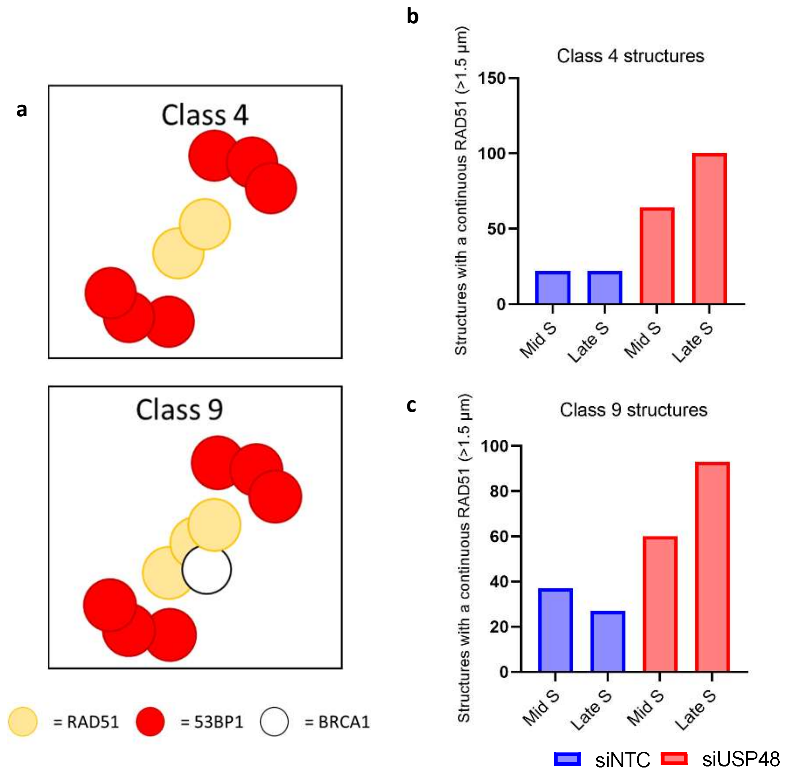


**Supplementary Figure 10. Prevalence of continuous RAD51 in specific RAD51, BRCA1 and 53BP1 enriched structures.**

a) Schematic representations of class 4 structures (defined as multiple RAD51 spots associated with multiple 53BP1 spots and no BRCA1 spots) and class 9 structures (defined as multiple RAD51 spots with BRCA1 spot(s) encapsulated by multiple 53BP1 spots) are shown.

b) The number of Class 4 structures with a continuous RAD51 structure (defined as continuous RAD51 of >1.5 µm, equivalent to >0.375 µm pre-expansion) were quantified from two independent experiments of control siRNA treated cells (siNTC) and USP48 siRNA treated cells. NTC mid S = 22 structures, NTC late S = 22 structures, siUSP48 mid S = 64 structures, siUSP48 late S = 100 structures.

c) The number of class 9 structures with a continuous RAD51 structure (defined as continuous RAD51 of >1.5 µm, equivalent to >0.375 µm pre-expansion) were quantified from two independent experiments of control siRNA treated cells (siNTC) and USP48 siRNA treated cells. NTC mid S = 37 structures, NTC late S = 27 structures, siUSP48 = 60 structures, siUSP48 = 93 structures.
